# Single-cell and spatial transcriptomics identify a macrophage population associated with skeletal muscle fibrosis

**DOI:** 10.1101/2023.04.18.537253

**Authors:** Gerald Coulis, Diego Jaime, Christian Guerrero-Juarez, Jenna M. Kastenschmidt, Philip K. Farahat, Quy Nguyen, Nicholas Pervolarakis, Katherine McLinden, Lauren Thurlow, Saba Movahedi, Jorge Duarte, Andrew Sorn, Elizabeth Montoya, Izza Mozaffar, Morgan Dragan, Shivashankar Othy, Trupti Joshi, Chetan P. Hans, Virginia Kimonis, Adam L. MacLean, Qing Nie, Lindsay M. Wallace, Scott Q. Harper, Tahseen Mozaffar, Marshall W. Hogarth, Surajit Bhattacharya, Jyoti K. Jaiswal, David R. Golann, Qi Su, Kai Kessenbrock, Michael Stec, Melissa J. Spencer, Jesse R. Zamudio, S. Armando Villalta

## Abstract

The monocytic/macrophage system is essential for skeletal muscle homeostasis, but its dysregulation contributes to the pathogenesis of muscle degenerative disorders. Despite our increasing knowledge of the role of macrophages in degenerative disease, it still remains unclear how macrophages contribute to muscle fibrosis. Here, we used single-cell transcriptomics to determine the molecular attributes of dystrophic and healthy muscle macrophages. We identified six novel clusters. Unexpectedly, none corresponded to traditional definitions of M1 or M2 macrophage activation. Rather, the predominant macrophage signature in dystrophic muscle was characterized by high expression of fibrotic factors, galectin-3 and spp1. Spatial transcriptomics and computational inferences of intercellular communication indicated that spp1 regulates stromal progenitor and macrophage interactions during muscular dystrophy. Galectin-3^+^ macrophages were chronically activated in dystrophic muscle and adoptive transfer assays showed that the galectin-3^+^ phenotype was the dominant molecular program induced within the dystrophic milieu. Histological examination of human muscle biopsies revealed that galectin-3^+^ macrophages were also elevated in multiple myopathies. These studies advance our understanding of macrophages in muscular dystrophy by defining the transcriptional programs induced in muscle macrophages, and reveal spp1 as a major regulator of macrophage and stromal progenitor interactions.

## INTRODUCTION

Macrophages have a central role in innate immunity and contribute to tissue homeostasis by regulating tissue repair and remodeling of the extracellular matrix (ECM) (*1*). Their diverse functions within the tissue are matched by a high degree of molecular heterogeneity, reflecting microenvironmental adaptability (*2–4*). Skeletal muscle macrophages comprise resident and monocyte-derived populations, the latter infiltrating upon muscle injury or disease (*5*). The M1 and M2 macrophage paradigm (*6, 7*) has been used to describe the functional heterogeneity of skeletal muscle macrophages (*8*). In acute muscle trauma, pro-inflammatory M1 macrophages initially infiltrate injured muscle to phagocytose cellular debris and activate muscle stem cells (*9–11*). The subsequent transition to M2 macrophages in the regenerative phase promotes muscle stem cell differentiation and the resolution of inflammation (*5, 12*). M1- and M2-like macrophages have been described in Duchenne muscular dystrophy (DMD) (*12, 13*). However, growing evidence indicates that the regulation and functional role of macrophages are more complex in the context of chronic muscle degenerative diseases such as DMD.

Although macrophages are essential for muscle repair following acute trauma, their dysregulation promotes the pathogenesis of DMD, a lethal form of muscular dystrophy caused by mutations in the *DMD* gene. The transition from M1 to M2 macrophages seen in acute injury is disrupted in the mdx mouse model of DMD by asynchronous bouts of muscle injury and regeneration (*14, 15*). Consequently, M1 macrophages are chronically activated and promote muscle injury in an iNOS-dependent manner (*4*), and the reparative function of M2 macrophages is pathologically repurposed to promote fibrosis (*12*). This impairment is expected to induce macrophage transcriptional programs that are distinct from those induced in acute injury and contribute to muscle pathology. In support of this, several studies reported that perturbing macrophage function and/or activation impaired regeneration and promoted fibrosis in muscular dystrophy (*16–19*) Pompe disease (*20*) and dysferlinopathy (*21*).

Fibrosis is the aberrant accumulation of collagen and other ECM proteins in chronically inflamed tissues, leading to organ failure and death (*22*). Fibro/adipogenic progenitors (FAPs) are stromal cells that give rise to fibroblasts and adipocytes, and regulate muscle repair and fibrosis (*23*). Following acute injury, FAPs expand and contribute to muscle repair by facilitating myogenesis and ECM formation (*24, 25*). Fibrosis is mitigated by infiltrating macrophages that clear FAPs through tumor necrosis factor alpha (TNFα)-mediated apoptosis (*26*). Conversely, TGF-β, which is highly upregulated in dystrophic muscle (*27, 28*), inhibits FAP apoptosis and guides their differentiation into matrix-producing myofibroblasts. In this setting, persistence of FAPs and their skewed differentiation towards fibroblasts contributes to the development of muscle fibrosis. Unexpectedly, inhibition of TGF-β does not fully restore FAP clearance (*26*), indicating that there are additional factors that promote FAP differentiation and/or survival. Osteopontin (spp1) is a potential candidate because it is highly expressed by macrophages (*19, 29*), it is elevated in dystrophic muscle in DMD patients (*30*) and promotes fibrosis in several disease settings (*31, 32*), including muscular dystrophy (*19, 30, 33*). However, the repertoire of pro-fibrotic factors, including spp1, expressed by dystrophic muscle macrophages and their cellular targets has not been fully defined.

In this study, we used an unbiased single-cell RNA sequencing (scRNAseq) approach to define the transcriptional profiles of macrophages from normal and dystrophic muscle. The scRNAseq identified several macrophage populations with novel transcriptomes not previously associated with muscular dystrophy. We focused on three populations that corresponded to resident macrophages, monocyte-derived macrophages, and a novel population characterized by high expression of pro-fibrotic factors, *Lgals3* (galectin-3) (*34–36*) and *Spp1* (*37, 38*). Given the selective induction of *Lgals3* and *Spp1* in dystrophic muscle macrophages, we hypothesize that this transcriptional profile defines a fibrogenic macrophage that promotes fibrosis during muscular dystrophy.

We demonstrate that galectin-3^+^ macrophages are activated in response to acute injury and are elevated in several muscle disorders. Although this macrophage population is transient following acute injury, galectin-3^+^ macrophages are chronically activated during muscular dystrophy. Spatial transcriptomic analysis of dystrophic muscle revealed that areas enriched in galectin-3^+^ macrophages and stromal cells expressed genes associated with muscle fibrosis. Furthermore, galectin-3^+^ macrophages colocalize with stromal cells in dystrophic lesions, and computational analysis with CellChat shows that Spp1 mediates communication between these cell types. Collectively, results from this study identify a distinct transcriptional profile in dystrophic muscle macrophages that is associated with fibrosis, revealing the potential for new therapeutic strategies that target fibrogenic macrophages in muscular dystrophy.

## RESULTS

### Single-cell RNA sequencing reveals novel cell states within skeletal muscle macrophages

Single-cell RNAseq was performed to unbiasedly phenotype skeletal muscle macrophage transcriptomes at single-cell resolution. We used a droplet-based scRNAseq platform (10X Genomics Chromium) to profile muscle macrophages (live F4/80^+^CD11b^+^siglec^-^ cells) purified by fluorescence-activated cell sorting (FACS) from mdx mice hind limbs during the acute stage of disease (4 weeks of age) and age-matched wildtype (WT) controls (**Fig. 1A**). FACS yielded macrophage samples with greater than 92-96% purity.

**Fig. 1.**
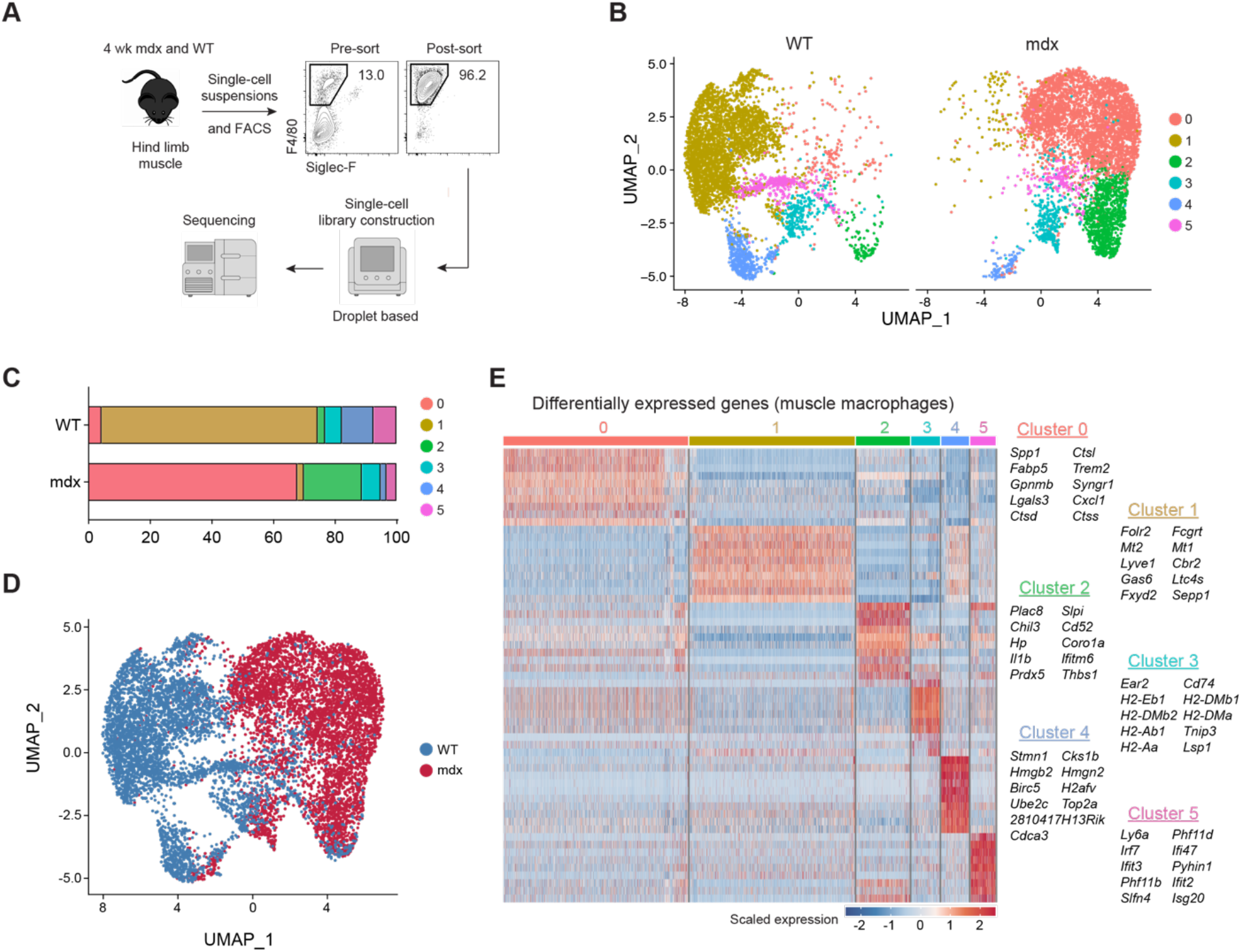
Identification of novel transcriptional programs in skeletal muscle macrophages by single-cell RNA sequencing. **(A)** Muscle macrophages were isolated from 4-wk-old WT (healthy) and mdx (dystrophic) mice and analyzed by scRNAseq. n= 1 **(B)** Dimensionality reduction via UMAP of healthy or dystrophic muscle macrophages. (**C**) The proportion of WT or mdx muscle macrophage clusters. (**D**) Identities classified by genotype. **(E)** Differential gene expression analysis showing the top ten most differentially expressed genes for each cluster.

Uniform Manifold Approximation and Projection (UMAP) for dimensionality reduction of 11,367 single-cell profiles (WT= 5723; mdx= 5644) was used to examine the data. The resulting analysis partitioned single-cell profiles into 8 clusters composed primarily of macrophages and a low proportion of contaminating cell types, including endothelial cells, FAPs, satellite cells and muscle-like cells (**fig. S1, A and B**). Clusters 0, 1, 2 and 3 made up more than 90% of single-cell profiles and were assigned a macrophage identity based on their expression of the macrophage markers *Cd68, Fcfr2b, Cd14* and *Adgre1* (**fig. S1C**). We filtered out the contaminating cell types and re-clustered the single-cell profiles corresponding to clusters 0-3 in fig. S1A, which resulted in the identification of 6 macrophage clusters (**fig. S1D**).

The proportion of clusters 0-5 macrophages differed among healthy and dystrophic muscle (**Fig. 1, B and C**). Cluster 1 macrophages were almost exclusively present in healthy muscle, whereas clusters 0 and 2 were predominantly found in dystrophic muscle. Clusters 3-5 were shared between healthy and dystrophic muscle. Vast differences in transcriptional profiles were noted between macrophages isolated from dystrophic (pink) and WT, healthy muscle (teal) (**Fig. 1D**). Hierarchical cluster analysis of differentially expressed genes revealed that each macrophage population expressed distinct transcriptional modules containing cluster-specific genes with biomarker potential (**Fig. 1E**).

Cluster 1 macrophages were characterized by increased expression of *Folr2, Mt2, Lyve1, Gas6* and *Cbr2*. This signature shared similarities with a subset of skeletal muscle-resident macrophages (SkMRMs) previously described in healthy muscle (*39*). Hereafter, cluster 1 macrophages are referred to as SkMRMs. Cluster 0 macrophages (hereafter referred to as galectin-3^+^ macrophages) expressed high levels of *Spp1, Fabp5, Gpnmb, Trem2, Lgals3* and various cathepsin genes. *Lgals3* (galectin-3), *Fabp5*, and *Gnmbp* have been implicated in the development of fibrosis in the heart, skin and liver (*40–43*), suggesting that galectin-3^+^ (gal-3^+^) macrophages promote fibrosis during muscular dystrophy. In support of this, *Spp1* (osteopontin) promotes muscle fibrosis in mdx mice (*30*) through a matrix metalloproteinase-mediated processing of TGFβ in stromal cells (*33*).

Cluster 2 macrophages, hereafter referred to as monocyte-derived macrophages (MDMs), were marked by *Cd52, Plac8, Prdx5* and *Hp*. F4/80 (*Adgre1*), which is lowly expressed in blood monocytes (*44*) (**fig. S2A**), was expressed lower in MDMs relative to SkMRMs or gal-3^+^ macrophages (**fig. S2B**). *Ly6c2* and *Cd52*, which are highly expressed in blood monocytes (**fig. S2, C and E**), were also highly expressed in MDMs, cluster 3 and 5 (**fig. S2, D and F**). Flow cytometry analysis of healthy and dystrophic muscle macrophages confirmed these observations (**fig. S2, G-J**). Further, MDMs expressed high levels of *Ccr2* but low levels of *Cx3cr1* (**fig. S2, K and L**). Collectively, these findings suggest that cluster 2 is a monocyte-derived population that resembles Ly6c^hi^CCR2^+^ inflammatory monocytes.

Cluster 3 was defined by high expression of *Cd74*, MHC II genes (*H2-Eb1, H2-DMb2, H2-Ab1, H2-Aa, H2-DMb1, H2-DMa*), and dendritic cells markers (*Itagx*), suggesting that this population corresponds to dendritic cells (*45*) (**Fig. 1E, fig. S2M**). Cluster 4 macrophages expressed genes associated with chromatin, nucleosome (*H2afv, Top2a, Hmgb2 and Hmgn2*) and cell cycle (*Birc5, Cks1b, Cdca3*) regulation. Cluster 5 defined a macrophage population characterized by the high expression of genes associated with interferon signaling (*Irf7, Ifi47, Ifit3, ifit2, Isg20*).

### Isolation and bulk transcriptome profiling of SkMRMs, monocyte-derived and galectin-3^+^ muscle macrophages by bulk RNAseq

A flow cytometry panel was developed to validate the scRNAseq macrophage populations that were predominantly associated with either homeostasis or dystrophinopathy. Galectin-3, Folr2 and CD52, which were preferentially expressed by clusters 0, 1 and 2, respectively, were used as markers to distinguish gal-3^+^ macrophages, SkMRMs and MDMs (**fig. S3, A and B**). Macrophages in WT muscle did not express galectin-3 but highly expressed Folr2, similar to SkMRMs identified in the scRNAseq analysis (**fig. S3, A and B**). In healthy tissues, Folr2^+^ macrophages were most abundant in skeletal muscle (72% ±2.5) followed by the heart (30% ±3.4) (**fig. S4**), but nearly absent in the brain, bone marrow and blood. This expression pattern expands on the observations made in an earlier study examining Folr2 in tissue-resident macrophages (*46*). A gal-3^hi^Folr2^lo^ macrophage population was also identified in dystrophic muscle that corresponded to gal-3^+^ macrophages in the scRNAseq (**fig. S3, A and B**), which was largely absent in several healthy tissues (**fig. S4**). CD52 was most highly expressed in gal-3^-^Folr2^-^ (WT) or gal-3^lo^Folr2^-^ (mdx) macrophages, and marked a population that likely corresponded to the MDMs in the scRNAseq analysis (**fig. S3, A and B**). CD52^+^ monocytes or macrophages were present in all WT tissues examined with the highest proportion in the blood, lung, liver and adipose tissue (**fig. S4**). Collectively, the flow cytometry panel discriminated three muscle macrophage populations based on galectin-3, Folr2 and CD52 expression.

To further link the flow cytometry macrophage populations to the scRNAseq profiles and establish their full transcriptomes, we performed bulk RNAseq on FACS-sorted populations. The macrophage populations were sorted from 4-wk-old WT (SkMRM) and dystrophic hind limb muscle (gal-3^hi^Folr2^lo^ and MDMs). The top 100 scRNAseq differentially expressed genes (scDEGs) from each cluster were used to match transcriptomes to the sorted populations. The SkMRM, gal-3^+^ and MDM scDEGs nearly uniformly distinguished the three transcriptomes and confirmed their match to the single cell populations (**Fig. 2, A-C; fig. S3, C-E**). Additionally, the gal-3^+^ scDEGs indicated partial overlapping profiles with the MDM scDEGs suggesting a cell state transition (**Fig. 2B**). Consistent with this, the gal-3^+^ population downregulated resident (*Folr2, Gas6, Cbr2, Lyve1*) and monocyte-derived macrophage (*Cd52, Ccr2, Ly6c2*) marker genes (**fig. S3F**). These observations suggest that the galectin-3^+^ population is a terminal transition state of either resident or monocyte-derived macrophages.

**Fig. 2.**
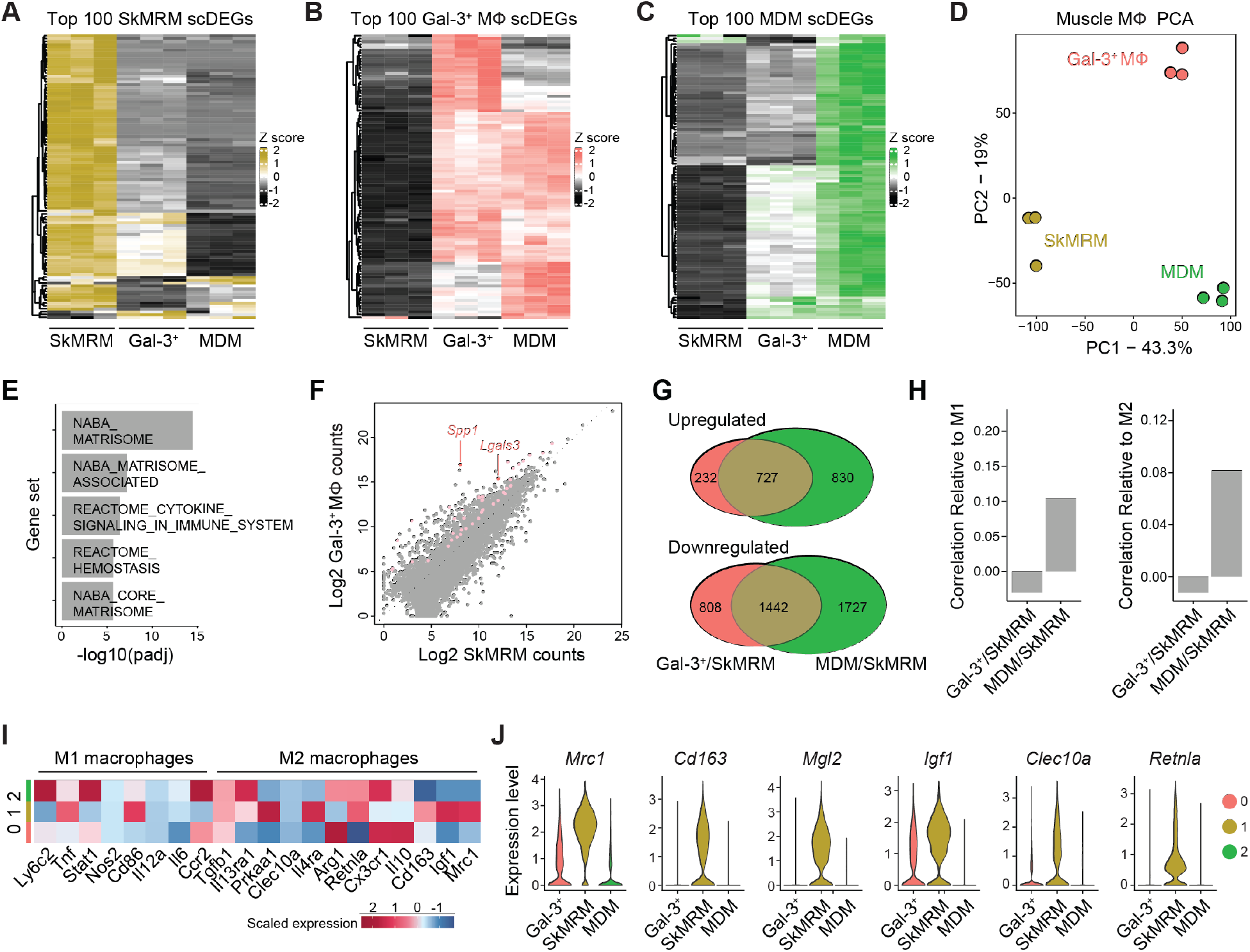
Muscle macrophages express novel transcriptomes, distinct from M1 and M2 macrophages. **(A-C)** Heatmap showing the expression of the top 100 scRNAseq differentially expressed genes from the scRNAseq analysis (scDEGs) in FACS-sorted SkMRM (A), gal-3^+^ M< (B) and MDM (C). n= 3 per population. (**D**) Principal component analysis (PCA) applied to the FACS-sorted macrophage populations in A-C. (**E**) Pathway analysis of top gene sets enriched in FACS-sorted gal-3^+^ M< compared to SkMRM. (**F**) Pairwise comparison of gene expression between FACS-sorted gal-3^+^ M< and SkMRM. Colored points indicate DEGs from ECM-related gene sets. *Lgals3* and *Spp1* are highlighted by arrows. (**G**) Venn Diagrams of upregulated and downregulated DEGs in FACS-sorted gal-3^+^ M< compared to SkMRM (Gal-3^+^/SkMRM) and MDMs compared to SkMRMs (MDMs/SkMRMs). (**H**) Correlation analysis of DEGs from Gal-3^+^/SkMRM or MDM/SkMRM comparisons with M1 or M2 polarized M<. (**I**) Expression of M1 and M2 macrophage markers in gal-3^+^ M< (0), SkMRMs (1) and MDMs (2) from scRNAseq data. (**J**) Violin plots of M2 markers in gal-3^+^ M<, SkMRMs and MDMs from scRNAseq data.

To further probe the macrophage transcriptomes, we performed principal component analysis (PCA) and classified DEGs. Using the SkMRM dataset and publicly available microglia and bone marrow-derived macrophage (BMDM<) datasets (*47, 48*), the analysis showed that ECM and development genes were enriched in SkMRMs, suggesting a role in muscle homeostasis (**fig. S5**). The PCA was next used to classify the variation between the sorted gal-3^hi^ macrophages, MDMs and SkMRMs (**Fig 2D**). The largest variance (PC1) separated normal and dystrophic macrophages and contained genes enriched in the innate immune response (**fig. S3G**). The second largest (PC2) variation was enriched for genes in pathways associated with the lysosome and lipid metabolism, suggesting that gal-3^+^ macrophages exhibit enhanced phagocytosis of muscle debris and lipid membranes (**fig. S3H**). By directly comparing DEGs between the gal-3^+^ macrophages and SkMRMs, ECM and inflammation-related pathways were further identified as highly altered by the dystrophic milieu (**Fig. 2E**). We determined 959 and 2250 upregulated and downregulated genes, respectively, in the gal-3^+^ macrophages compared to SkMRMs (**fig. S3I, table S1**). Consistent with marker gene upregulation, *Lgals3* and *Spp1* were among the highest upregulated and expressed genes in the galectin-3^+^ state (**Fig. 2F**).

The common transcriptional response of gal-3^+^ macrophages and MDMs to the dystrophic environment was indicated by an overlap of 727 upregulated (41% of total) and 1442 downregulated (36% of total) DEGs compared to SkMRMs (**Fig. 2G**, p-values > 1e-16). Pathways enriched within the common DEGs indicated upregulated genes involved in leukocyte activation and the inflammatory response, and downregulated genes in glycosaminoglycan and ECM-related pathways (**Fig S3J**). Gal-3^+^ macrophages and MDMs also expressed unique DEGs, indicating specialized functional states for distinct dystrophic muscle macrophage populations. The analysis of enriched pathways in the unique DEGs again indicated ECM and lysosomal gene upregulation in the galectin-3^+^ state compared to the MDMs, potentially reflecting enhanced fibrotic and phagocytic activity (**fig. S3, K and L**).

To further characterize the inflammatory states of the dystrophic macrophages, we compared them to well-defined M1 and M2 polarization signatures established by *in vitro* treatments of BMDM< (*48*). The transcriptional profiles substantially differed between muscle and BMDM< macrophages (**fig. S3M**), and little to no correlation was found between the MDM (C2/C1) or galectin-3^+^ state (C0/C1) and M1 or M2 polarization states (**Fig. 2H**). Further, several M1 and M2 markers were heterogeneously expressed across the dystrophic macrophage populations in the scRNAseq analysis (**Fig. 2I**). Unexpectedly, some M2 markers were expressed highest in SkMRMs (**Fig. 2J**). Flow cytometry and qPCR analysis confirmed that M1 and M2 genes were not selectively enriched in any of the scRNAseq-defined muscle macrophage populations (**fig. S6**). The transcriptomic profiling of muscle macrophages supports that a terminal differentiation state marked by high galectin-3 expression is a dominant signature induced in muscular dystrophy. This galectin-3 signature is associated with regulation of the ECM, and has little overlap with complete M1 and M2 macrophage activation signatures.

### Spatial transcriptomic analysis reveals that galectin-3^+^ macrophages are associated with stromal cells and ECM genes

We used capture probe-based spatial transcriptomics (Visium, 10X Genomics) to gain insight on how gal-3^+^ macrophages functionally and spatially interfaced with the dystrophic environment. Spatial RNA sequencing was performed on the gastrocnemius/plantaris muscle complex of 6-wk-old mdx mice in the DBA2/J background (D2-mdx). This approach provides spatially-resolved gene expression analysis limited in resolution by 55 μm diameter spot size. Interrogation of the spatial matrix revealed that areas with high galectin-3 expression (**Fig. 3A**, highlighted in red) were confined to regions with active pathology that were densely populated by mononuclear cells (**Fig. 3B**, highlighted in yellow).

**Fig. 3.**
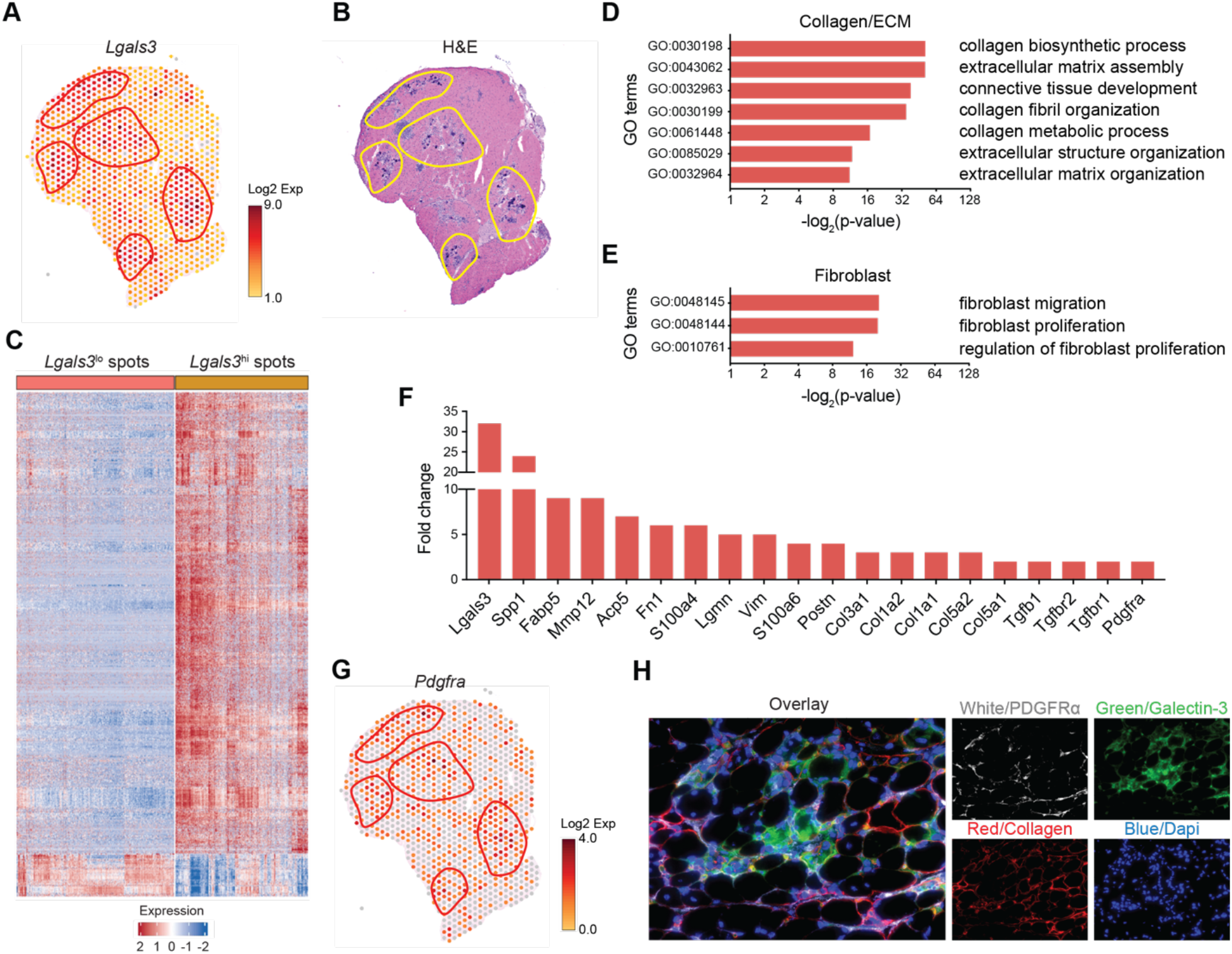
Spatial transcriptomics reveals that Gal-3^+^ macrophages are associated with stromal cells and extracellular matrix. (**A** and **B**) Spatially resolved gene expression of *Lgals3* (galectin-3) (A) and H&E staining of D2-mdx quadriceps (B). Shown is one of 5 representative D2-mdx quadriceps. (**C**) Heatmap showing DEGs between Lgals3^hi^ and Lgals3^lo^ spots. Shown are genes with a fold change >= 1.5, FDR < 0.01. All spots in a section, including those with and without pathology, were unbiasedly analyzed. (**D** and **E**) Gene ontology/pathway analysis showing the enrichment of GO terms associated with Collagen/ECM (D) and fibroblasts (E) in galectin-3^hi^ spots. (**F)** Expression of DEGs associated with fibrosis. (**G**) Spatially resolved gene expression of *Pdgfra* in mdx quadriceps. (**H**) Immunofluorescence staining of 4-wk-old mdx quadriceps reveals colocalization of PDGFRa and galectin-3 in dystrophic muscle.

A differential gene expression analysis between galectin-3^hi^ and galectin-3^lo^ spots revealed that 1,365 and 125 genes were up and downregulated, respectively (**Fig. 3C**). The provided heatmap comprises 569 galectin-3^hi^ and 676 galectin-3^lo^ spots collected from five D2-mdx gastrocnemius/plantaris muscle complexes. We assigned any spot with a UMI count ≥ 3 as galectin-3^hi^, and spots with a UMI count ≤ 1 as galectin-3^lo^. A gene ontology (GO) analysis was performed on the DEGs, which showed that phagocytosis, endocytosis and leukocyte migration were among the most enriched terms (**Fig. S7A**). Terms associated with regeneration and repair were also highly enriched in the galectin-3^hi^ areas (**Fig. S7B**). Of particular interest, GO terms associated with the ECM and fibroblasts (**Fig. 3, D and E**), and multiple genes associated with fibrosis, were enriched in the galectin-3^hi^ areas (**Fig. 3F**). *Lgals3* was among the highest, revealing a greater than 30-fold increase. Consistent with our scRNAseq analysis, showing that gal-3^+^ macrophages also express S*pp1*, we found that galectin-3^hi^ areas expressed high levels of *Spp1* (∼25-fold increase). The pro-fibrotic matrix metallopeptidase-12 (*Mmp12*) was also increased (*49*). Genes encoding ECM components were also increased, including fibronectin (*Fn1*), periostin (*Postn*), and several collagens, as well as components of growth factor pathways that induce fibrosis (e.g. TGFβ and PDGF).

The spatial transcriptomics also revealed that PDGFRα, a growth factor receptor expressed on stromal cells, was increased in the galectin-3^+^ areas (**Fig. 3G**, highlighted in red). Immunofluorescence assays performed on mdx quadriceps revealed that PDGFRα^+^ stromal cells in pathological lesions were juxtaposed with galectin-3^+^ macrophages (**Fig. 3H**). Collectively, these findings suggest that galectin-3^+^ macrophages interact with stromal cells (e.g. FAPs) in degenerative lesions to promote fibrosis.

### Intercellular communication network analysis identifies that FAPs and macrophages communicate via spp1

To determine how gal-3^+^ macrophages and stromal progenitors (i.e. FAPs) interact, we performed a reference-based integration of skeletal muscle mononucleated cell datasets and surveyed intercellular communication networks with CellChat (*50*). We used a publicly available dataset of uninjured muscle (*51*) and referenced a dataset prepared from muscle mononucleated cells pooled from 3 mdx mice. Three subpopulations of FAPs, including adipogenic, pro-remodeling and stem populations were identified, as reported previously (*52*), in uninjured and dystrophic muscle. An increased proportion of pro-remodeling (22.8% vs 8.6%) and stem FAPs (20.5% vs 10.8%) was noted in dystrophic muscle, as well as an overall expansion of muscle macrophages (38.9% vs 14.4%), compared to healthy muscle (**Fig. 4A**). Similar to the scRNAseq analysis of purified macrophages (**Fig. 1**), gal-3^+^ macrophages were the predominant macrophage population in dystrophic muscle (**Fig. 4A**). The probability of communication between two cell groups was visualized with circle plots by designating a FAP population as the central nodes of analysis in the mdx dataset (**Fig. 4, B-D**). Although a complex communication network was present for all FAPs, the highest degree of inferred communication occurred with gal-3^+^ macrophages (reflected by the thickness of the connecting edge).

**Fig. 4.**
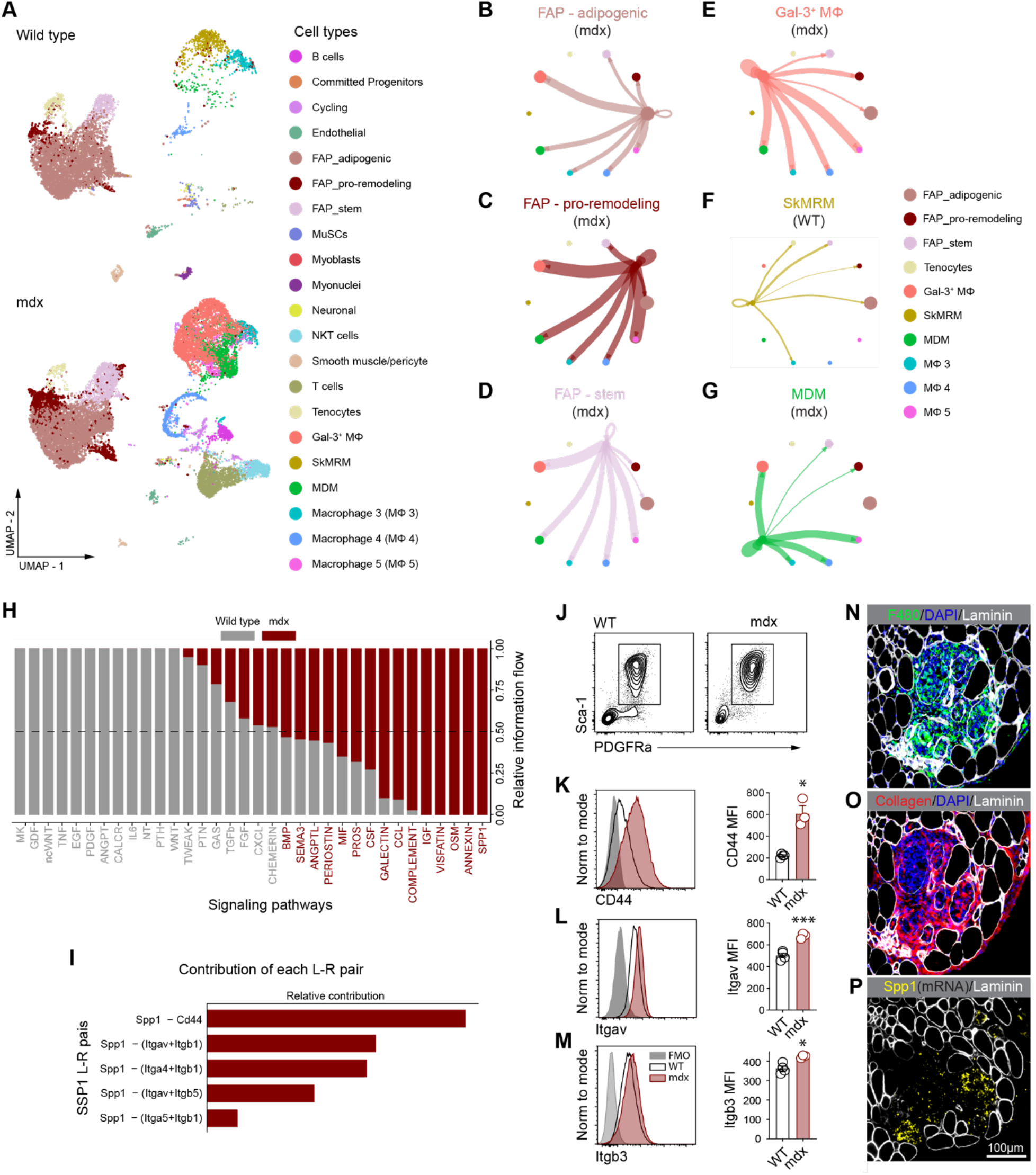
Spp1 mediates FAP and macrophage interactions in dystrophic muscle. **(A)** reference-based integration of skeletal muscle mononucleated cell datasets prepared from 3-mon-old WT and mdx mice. (**B-G**) Visualization and analysis of cell-cell communication using CellChat. Circle plots placing FAPs subsets as the central nodes of analysis in the mdx dataset (B-D). An interaction between a pair of cell types is depicted by a line connecting two cell types. The thickness of the line depicts the strength of the interaction between two cell types. A similar analysis applying macrophages as the central node of communication is shown (E-G). (**H**) Pathways enriched in the stromal cell and macrophage network of WT and mdx mice. (**I**) Relative contribution of Spp1 ligand (L)-receptor (R) pairs. (**J**) Expression of Spp1 receptors was measured in WT and mdx PDGFRα^+^Sca1^+^ FAPs by flow cytometry. Plots shown were gated on CD45^-^CD31^-^ live cells. (**K-M**) Representative histograms and quantification of the MFI of Spp1 receptors. 4-wk-old mice were analyzed. n= 3-4. *p<0.05, **p<0.01, ***p<0.001 using an unpaired Welch’s t-test. (**N-P**) RNAscope multiplexed with immunofluorescence staining showing the co-localization of macrophages (N) and Spp1 mRNA (P) in areas enriched with collagen (O).

The CellChat analysis was also performed with gal-3^+^ macrophages, SkMRM or MDMs as the central nodes of analysis (**Fig. 4, E-G**). Circle plots show a large probability of communication between gal-3^+^ macrophages and other macrophage subpopulations, followed by communication with pro-remodeling FAPs in the mdx dataset (**Fig. 4E**). MDMs principally communicated with other macrophage subpopulations and made negligible interactions with FAP subpopulations in mdx mice (**Fig. 4G**). Intercellular communication between tenocytes and macrophages was not seen in mdx mice. Interestingly, SkMRMs poorly communicated with other macrophage and FAP populations in uninjured muscle, suggesting that the function of SkMRMs differs from that of gal-3^+^ macrophages (**Fig. 4F**).

Next, information flow for all significant signaling pathways in the stromal cell and macrophage network was assessed (**Fig. 4H**). Some pathways (e.g. CXCL, CHEMERIN, BMP, SEMA3, ANGPTL, PERIOSTIN) maintain similar flow in healthy and dystrophic muscle, suggesting that these pathways are equally important in regulating macrophage and FAP interaction during homeostasis and muscle disease. In contrast, other pathways prominently change their information flow in dystrophic muscle compared to uninjured muscle. For example, MK and GDF are silenced in dystrophic muscle, and TWEAK, PTN and GAS are decreased. In contrast, GALECTIN, CCL and COMPLEMENT are increased in dystrophic muscle. IGF, VISFATIN, OSM, ANNEXIN and SPP1 were exclusively active in dystrophic muscle.

The Spp1 pathway was further characterized given its known role in promoting muscle fibrosis during muscular dystrophy (*30*). The highest relative contribution to the Spp1 pathway was attributed to the Spp1-CD44 ligand (L)-receptor (R) pair, followed by several integrin heterodimer receptors (**Fig. 4I**). *Cd44* was expressed highest in pro-remodeling FAPs, but was not detected in stem or adipogenic FAPs in mdx mice (**Fig. S8**), whereas integrins were heterogeneously expressed in all FAP populations. CD44 and integrins were also expressed in macrophage populations. The expression of Spp1 receptors in WT and mdx PDGFRα^+^Sca1^+^ stromal cells was measured by flow cytometry (**Fig. 4J**). CD44, Itgav and Itgb3 were increased in mdx stromal cells (**Fig. 4, K-M**). Consistent with the observation that PDGFRα^+^ stromal cells interacted with gal-3^+^ macrophages (Fig. 3H), RNAscope multiplexed with immunofluorescence staining revealed that degenerative lesions with increased macrophages and collagen contained elevated levels of *Spp1* (**Fig. 4, N-P**). Collectively, these results suggest that the Spp1 pathway is a key signaling axis controlling gal-3^+^ macrophage and FAP interactions during muscular dystrophy, and that Spp1 signals primarily through CD44 and a subset of integrin heterodimers.

### Muscle damage expands a population of galectin-3^+^ macrophages that is chronically activated in muscular dystrophy

We next examined the regulation of muscle macrophage populations during muscular dystrophy. SkMRM and MDMs were elevated in the early stages of disease but resolved by chronic stages (**fig. S9, A and B**). An elevation of gal-3^+^ (galectin-3^hi^Folr2^lo^) macrophages was observed as early as 3.5 weeks of age in mdx hind limb muscles and began to decline by 8 weeks, but remained chronically elevated up to 52 weeks of age compared to controls (**Fig. 5A**). Galectin-3 protein was upregulated in dystrophic muscle macrophages as early as 4 weeks and remained elevated at 52 weeks of age (**Fig. 5, B and C**). Immunofluorescence staining showed that galectin-3 was expressed in a subset of F4/80^+^ macrophages in dystrophic muscle, but absent in WT muscle macrophages (**Fig. S8C**). The chronic activation of gal-3^+^ macrophages was associated with increased collagen deposition at 52 weeks (**Fig. 5D**).

**Fig. 5.**
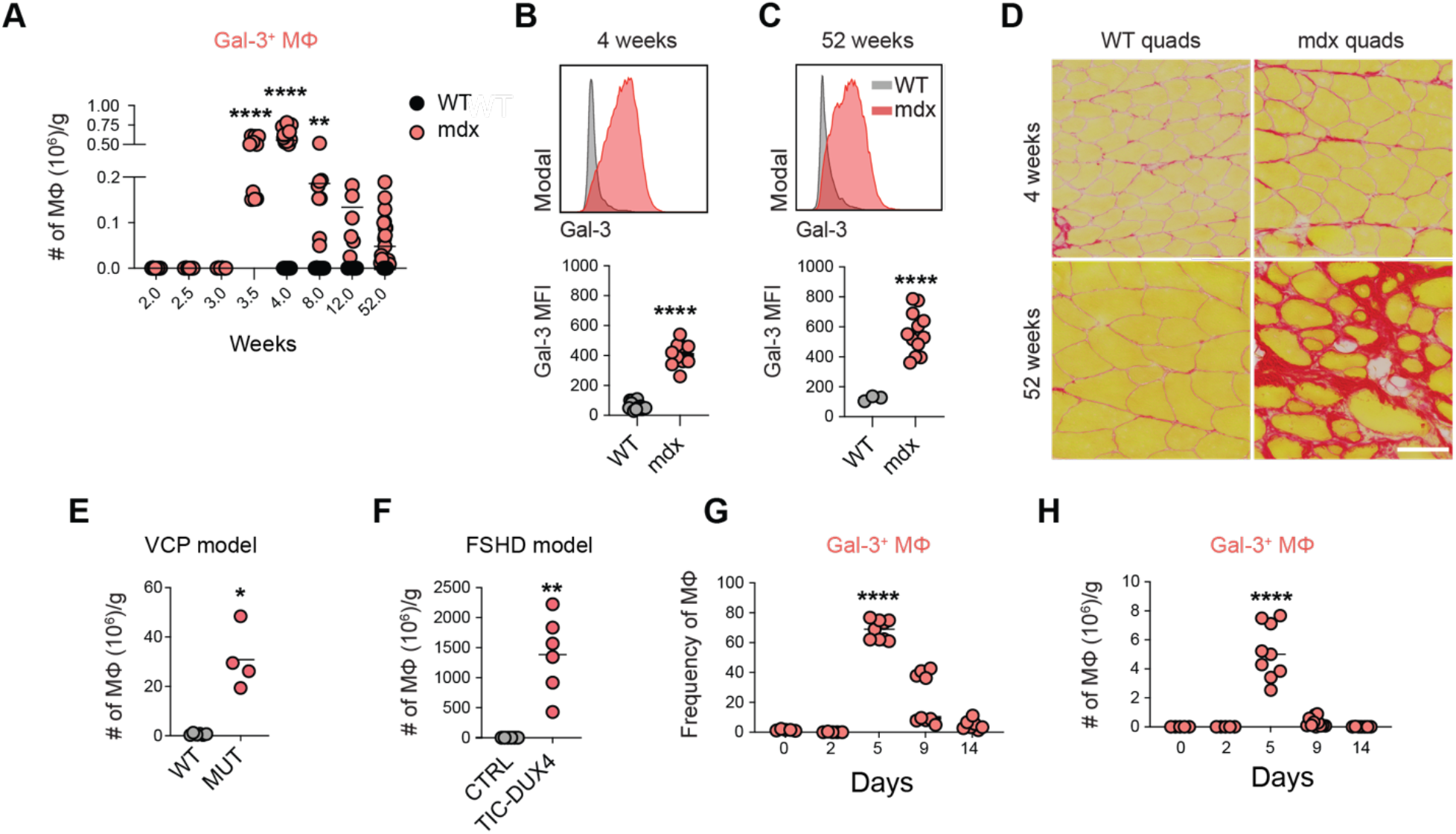
Chronic activation of galectin-3^+^ macrophages in dystrophic muscle. **(A)** The number of gal-3^+^ macrophages in B10.mdx hind limb muscle, normalized to muscle mass (g). n= 6-19 per time point (**B** and **C**) Representative histograms and quantitative analysis of the geometric mean fluorescence intensity (MFI) of Gal-3 in 4 wk-(B) and 52-wk-old (C) WT and mdx mice. n= 3-16 per group. 4-wk-old mice were analyzed. (**D**) Sirius red staining of WT and mdx quadriceps cryosections from 4 wk- and 52-wk-old mice. (**E-F**) Enumeration of gal-3^+^ macrophages in the valosin-containing protein (VCP)-associated inclusion body myopathy mouse model (E) and in the Facioscapulohumeral Muscular Dystrophy (TIC-DUX4) mouse model (F). n= 4-6, 10-mon-old mice (E); n= 5, 10-w-kold mice (F). (**G** and **H**) The regulation of gal-3^+^ macrophages frequency (G) and number (H) after injury. n= 7-9 per time point (G, H). *p<0.05, **p<0.01, ***p<0.001, ****p<0.0001 using an unpaired Welch’s t-test (B, C, E, F) or 2-way ANOVA with Sidak’s multiple comparison test (A, G, H).

The induction of a gal-3^+^ transcriptional program was conserved in other forms of muscle disease. We performed a referenced-based integration of muscle macrophage scRNAseq datasets prepared from 8-month-old B6A/J mice, a mouse model of limb girdle muscular dystrophy 2B, and respective controls (**fig. S10A**). The mdx dataset from this study (Fig. 1) was used as the reference. Similar to mdx dystrophic muscle, the predominant muscle macrophage population in B6A/J mice corresponded to gal-3^+^ macrophages (**fig. S10B**). Immunofluorescence staining of 12-month-old muscle confirmed the presence of gal-3^+^ macrophages and their proximity to PDGFRα^+^ stromal cells in B6A/J mice (**fig. S10D**). Gal-3^+^ macrophages were largely absent in B6 healthy control muscle (**fig. S10C**). SkMRMs were similarly the dominant macrophage population in control muscle. Although the lack of MDMs and cluster 5 macrophages in B6A/J and control samples could be explained by disease-specific differences, we cannot rule out that differences in the stage of disease (i.e. age) could contribute to this observation. Gal-3^+^ muscle macrophages were also elevated in mouse models of valosin-containing protein (VCP)-associated inclusion body myopathy (*53*) and Facioscapulohumeral Muscular Dystrophy (FSHD) (*54*) (**Fig. 5, E and F**).

The regulation of gal-3^+^ macrophages following BaCl_2_-induced acute injury was also examined. A large increase in the proportion and number of gal-3^+^ muscle macrophages occurred five days after injury (**Fig. 5, G and H**). By day 9, gal-3^+^ macrophages began to contract and returned to their homeostatic levels by day 14. The expansion of gal-3^+^ macrophages at day 5 coincided with the transition of muscle towards the repair phase and when collagen deposition was most apparent, suggesting that they are involved in remodeling of the extracellular matrix during muscle regeneration (**Fig. S11, A and B**).

### Galectin-3^+^ macrophages are derived from skeletal muscle-resident macrophages and peripheral monocytes

The capacity of SkMRMs and monocytes to differentiate into galectin-3^hi^Folr2^lo^ macrophages was determined through adoptive transfer assays. SkMRMs or bone marrow monocytes from CD45.1^+^ congenic WT mice were injected into the quadriceps of 4-wk-old mdx.CD45.2^+^ recipient mice. Prior to the adoptive transfer (day 0), bone marrow monocytes did not express Folr2 or galectin-3 (**Fig. 6A**). CD45.1^+^ transferred monocytes upregulated galectin-3 and Folr2, leading to an increased proportion of cells that acquired the gal-3^hi^Folr2^lo^ phenotype as early as day 2 and remained elevated 7 days after transfer (**Fig. 6, A and B**). The adoptive transfer of CD45.1^+^ SkMRMs from WT muscle resulted in a downregulation of Folr2 and an upregulation of galectin-3, resembling the galectin-3^hi^Folr2^lo^ phenotype by the second day after transfer (**Fig. 6, B and D**). CD45.1^+^ SkMRMs were substantially declined 7 days after transfer, suggesting that galectin-3^hi^Folr2^lo^ emerging from the SkMRM pool are short-lived (**Fig. 6D**).

**Fig. 6.**
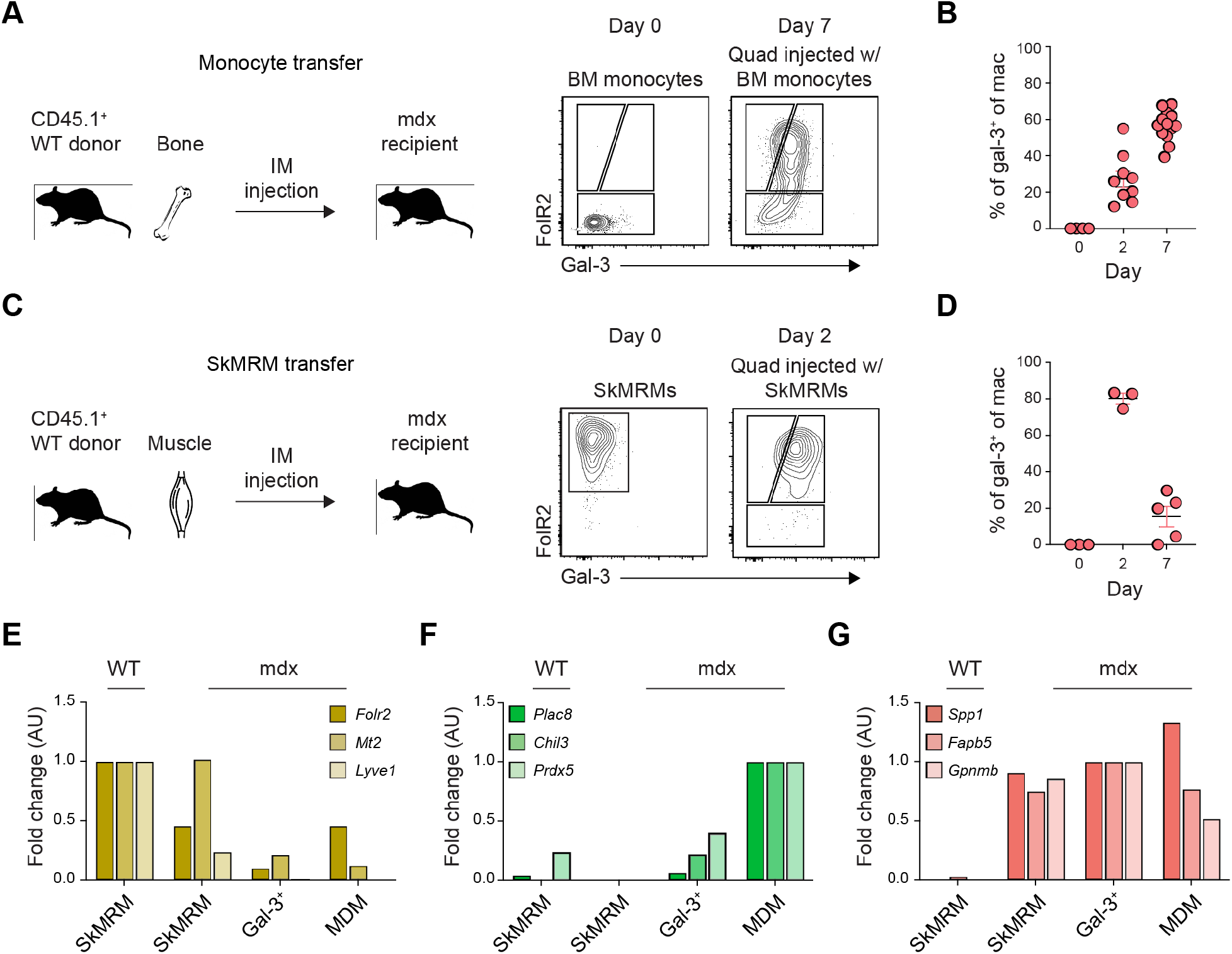
Peripheral monocytes and skeletal muscle-resident macrophages give rise to galectin-3^+^ macrophages. (**A** and **B**) Adoptive transfer of monocytes into 4-wk-old mdx mice. Graphical abstract of the workflow and representative flow plots of monocytes before and after transfer (A). Frequency of the donor monocytes that converted to gal-3^+^ macrophages at 2- and 7-days post-transfer (B). n= 3-13 per group. (**C** and **D**) Adoptive transfer of SkMRMs into 4-wk-old mdx mice. Schematic of the workflow and representative flow plots (C). Frequency of SkMRMs that converted to gal-3^+^ macrophages (D). n= 3-5 per group. (**E-G**) RT-qPCR quantification of the expression of cluster 1 (E), 2 (F) and 0 genes (G) in FACS-sorted SkMRMs from WT and mdx muscle, and *p<0.05, **p<0.01, ***p<0.001, ****p<0.0001 using an unpaired Welch’s t-test (B, C, E, F) or 2-way ANOVA with Sidak’s multiple comparison test (A, G, H). gal-3^+^ M< and MDMs from mdx muscle.

The regulation of SkMRMs, MDMs and gal-3^+^ macrophage marker genes in endogenous macrophages FACS-sorted from 4-wk-old WT and mdx muscle was assessed by RT-qPCR (**fig. 6, E-G**). The following marker genes were interrogated, *spp1, Fabp5* and *Gpnmb*; *Folr2, Mt2* and *Lyve1*; and *Plac8, Chil3* and *Prdx5*, because of their preferential expression in gal-3^+^ macrophages, SkMRMs and MDMs, respectively (**fig. S12**). Folr2^hi^ macrophages isolated from dystrophic muscle (mdx SkMRM) began to lose expression of SkMRM genes (**Fig. 6E**), but gained expression of gal-3^+^ macrophage genes (**Fig. 6G**). Similarly, MDMs in dystrophic muscle upregulated gal-3^+^ macrophage genes (**Fig. 6G**), whereas MDM genes were lowly expressed in gal-3^+^ macrophages (**Fig. 6F**). This regulation of SkMRM, MDM and gal-3^+^ macrophage genes is consistent with the adoptive transfer assays and supports the interpretation that SkMRM and recruited CD52^+^ monocytes are activated and differentiate into galectin-3^hi^Folr2^lo^ macrophage within the dystrophic milieu.

### Galectin-3^+^ macrophages are elevated in human diseased muscle and interact with FAPs

To determine if gal-3^+^ macrophages are also present in human dystrophic muscle, we quantified their numbers in archived muscle biopsies. Immunofluorescence assays showed that galectin-3 was expressed in a subset of CD68^+^ macrophages (**Fig. 7A**). An immunohistochemical examination of galectin-3 showed that the number of gal-3^+^ macrophages in interstitial or perivascular regions did not differ between control and myopathic patients, except for an increase in gal-3^+^ perivascular macrophages in limb girdle muscular dystrophy 2A (LGMD2A) (**Fig. 7, B-D**). However, the number of gal-3^+^ myofiber-invading macrophages was significantly elevated in DMD, antisynthetase syndrome (ASS) and LGMD2A (**Fig. 7, B and E**). A non-significant trend for an increase in inclusion body myositis (IBM) patients was also noted. Consistent with the increase in gal-3^+^ macrophages in DMD and LGMD2A, the expression of *Spp1* in whole muscle was elevated (**Fig. 7F**). RNA was not available for IBM, NAM and ASS groups to measure transcript levels of *Spp1*. Further, *COL1A* mRNA was significantly elevated in DMD muscle, suggesting that a similar gal-3^+^ macrophage and Spp1 pathway promotes fibrosis in DMD. (**Fig. 7, F and G**).

**Fig. 7.**
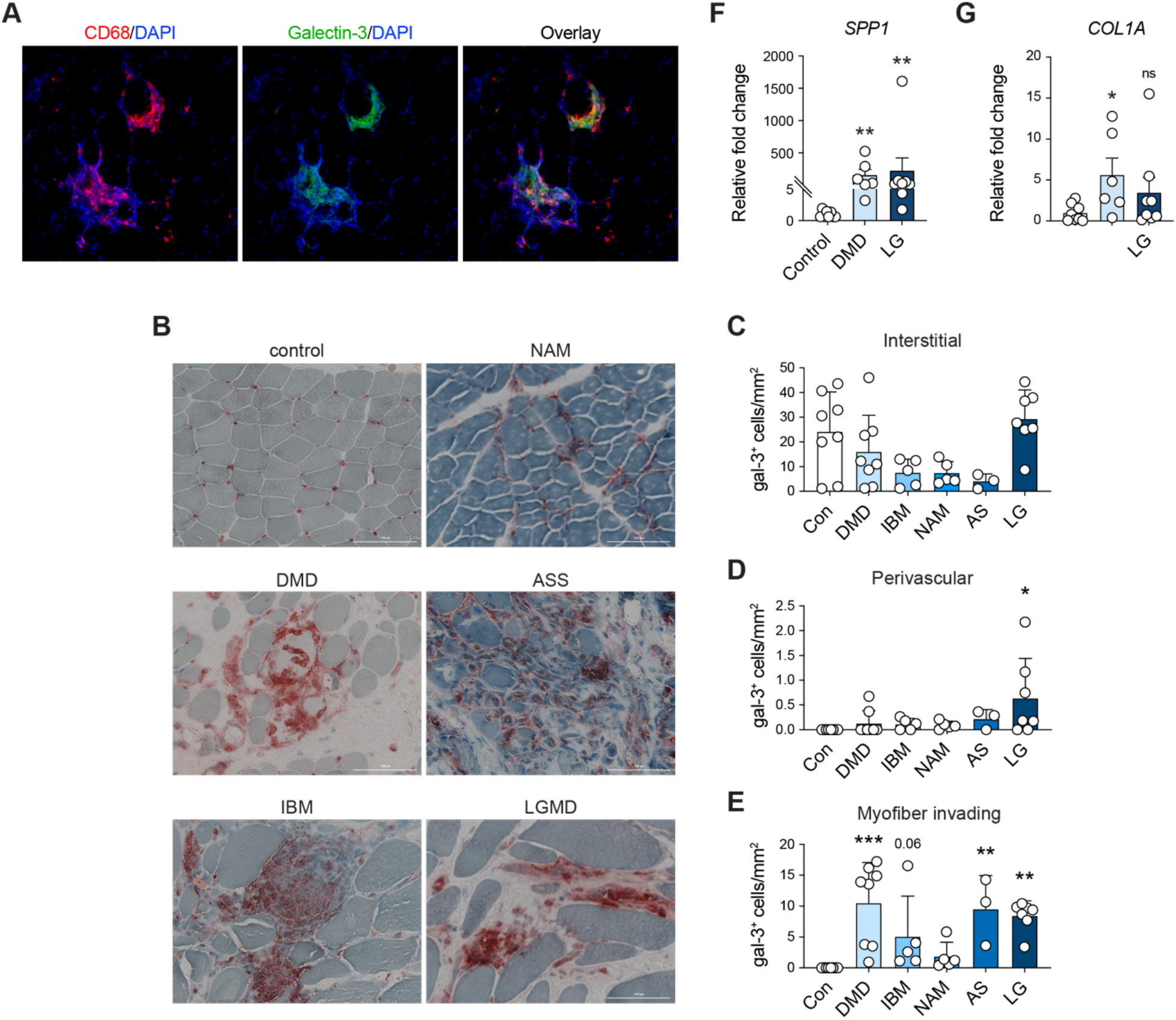
Galectin-3^+^ macrophages are elevated in human chronic muscle disease. (**A**) Immunofluorescence staining of gal-3^+^ macrophages in human dystrophic muscle. CD68 (red), gal-3 (green), nuclei (blue). (**B**) Representative images of immunohistochemical staining of galectin-3 in control and myopathic patients. (**C-E**) Quantification of gal-3^+^ cells in the interstitial space (C), the perivascular area (D) or infiltrating the myofiber (E). n= 3-8 frozen sections per patient type. (**F** and **G**) Expression of human SPP1 (F) and COL1A (G) mRNA in control, DMD and LGMD biopsies. n= 6-8 patients were used to measure RNA expression. *p<0.05, **p<0.01, ***p<0.001, ****p<0.0001 using a 2-way ANOVA with Kruskal-Wallis multiple comparison test (C-G).

## DISCUSSION

Macrophage diversity is regulated by the integration of tissue-intrinsic cues with inflammatory signals to induce unique transcriptional programs. Single-cell transcriptomics has helped uncover an unprecedented degree of macrophage heterogeneity attributed to the integration of these complex signals in several organs, including the heart, brain and adipose tissue (*55–60*). Recently, comparative single-cell transcriptomic studies of tissue-resident macrophages isolated from distinct anatomical sites, including skeletal muscle, demonstrated that muscle macrophages expressed a transcriptome that differs from those of lung and peritoneal macrophages (*39*). Transcriptomic analysis of muscle macrophages isolated from acutely-injured muscle has also shown that macrophages acquire distinct transcriptional programs adapted to promoting muscle regeneration (*10, 51, 52, 61*). However, the diversity of muscle macrophages, and how their function and transcriptional states are influenced by muscle disease, have not been defined.

Here, we performed an unbiased scRNAseq analysis of muscle macrophages from healthy and dystrophic muscle to define the transcriptional programs induced in macrophages that promote muscle fibrosis during muscular dystrophy. We found six distinct macrophage populations, none of which resembled polarized M1 or M2 macrophages. Rather, the six populations expressed varying degrees of M1 and M2 macrophage markers genes. The gal-3^hi^Folr2^lo^ (gal-3^+^) signature emerged as the predominant transcriptional state in dystrophic muscle macrophage. Gal-3^+^ macrophages expressed pro-fibrotic factors, including spp1 and galectin-3, and were chronically activated in muscular dystrophy. Gal-3^+^ macrophages interacted with stromal cells (e.g FAPs) in regions enriched with genes associated with muscle fibrosis, and computational inferences predicted that communication between these cells is partly mediated by Spp1. Gal-3^+^ macrophages were also elevated in several human myopathies, including DMD, and were juxtaposed with PDGFRα^+^ stromal cells. Further, spp1 and collagen were upregulated in DMD biopsies, suggesting that a spp1-mediated gal-3^+^ macrophage and stromal cell interaction is conserved in human to promote muscle fibrosis.

Consistent with a potential role for gal-3^+^ macrophage-derived Spp1 in the pathogenesis of muscular dystrophy, Spp1 enhances the matrix metalloproteinase 9-mediated processing of TGFβ into its active form to promote muscle fibrosis in mdx mice (*19, 30, 33*). Further, Spp1 skews macrophages towards a pro-inflammatory phenotype, which promote muscle damage in acute stages of disease in mdx mice (*13, 19*). Spp1 and galectin-3 are also associated with fibrosis in the heart, lung and kidney (*62*) (*63*) (*64*) (*34, 65, 66*), and adipogenic differentiation of FAPs in skeletal muscle (*67*), suggesting a cooperative role for galectin-3 and Spp1 in inducing fatty fibrosis in DMD. In support of this, Spp1 and galectin-3 induce fibroblast proliferation and differentiation of myofibroblasts (*68, 69*). It is essential to note, however, that galectin-3 may have a potential role in tissue repair as demonstrated in the brain (*70*) and lung (*71*). The emergence of gal-3^+^ macrophages has also been reported in the brain, lung, heart, aorta and liver when homeostasis is disrupted in these organs (*65, 72–76*). Studies of barium chloride-induced muscle injury or autologous muscle grafting in mice showed that galectin-3 and Spp1 promote regeneration in acute settings (*77–79*). These findings suggest that muscle regeneration and fibrosis share common pathways of regulation, and a temporal-dependent mechanism partly determines whether Spp1 and galectin-3 promote regeneration or fibrosis. This perspective is consistent with the view that fibrosis, to some extent, reflects a dysregulated repair process (*12, 80*). Thus, therapeutics that target a single inflammatory mediator (e.g. Spp1 or galectin-3) to inhibit fibrosis will likely be suboptimal approaches for complex disorders like muscular dystrophy, because of their dual role in regeneration. Rather, more effective strategies would center on inhibiting the transcriptional program associated with gal-3^+^ macrophages and reverting it to a homeostatic state.

In this regard, the present study defines the homeostatic signature of SkMRMs and establishes *a posteriori* knowledge required to further investigate how this transcriptional state is regulated. The transcriptional profile of SkMRMs identified in this study overlapped with resident macrophages identified by scRNAseq analysis of healthy skeletal muscle (*39*). Wang and colleagues described four macrophage clusters, including cluster 0, proliferating, Ccr2^+^ and Cd209^+^ clusters, the latter of which was enriched in healthy quadriceps. A resident population in the diaphragm was also characterized by high expression of *Folr2, Lyve1, Ltc4s, Fxyd2* and *Fcgrt*, genes that were expressed the highest in SkMRMs in the present studies. Unexpectedly, we found that markers associated with M2 macrophages were most highly expressed in SkMRMs. This observation was consistent with the high expression of M2 markers in the CD209^+^ muscle-resident macrophages described by the Wang et al. (*39*). Although SkMRMs expressed some marker of M2 macrophages, their overall transcriptional signature substantially differed from M2 macrophage polarized *in vitro* with IL-4 and IL-13. Given that the cytokines that induce M2 activation (e.g. IL-4 and IL-13) are increased with muscle injury or disease, and are typically low or absent in healthy muscle (*81, 82*), it is unlikely that these factors promote the M2-like phenotype of SkMRMs. Rather, tissue-associated or metabolic signals are more likely factors to induce this phenotype (*83*). Consistent with this interpretation, bulk RNAseq analysis revealed that SkMRMs were associated with ECM and muscle-associated pathways, suggesting that these homeostatic functions induce a molecular phenotype sharing some features with M2 macrophages.

A significant advancement in this study was the observation that the dystrophic environment converts muscle-resident macrophages and peripheral monocytes into gal-3^+^ macrophages. Prior studies implicated a role for recruited monocytes in the muscle pathology of mdx mice at 12 weeks of age (*15*). Although Ly6c^hi^ monocytes were reduced out to 6 months of age in CCR2-deficient dystrophic mice, myonecrosis and fibrosis returned to control levels by this point (*84*), suggesting that other monocyte or macrophage populations promote dystrophinopathy at later stages of disease. In this regard, Zhou and colleagues concluded that Ly6c^lo^ monocytes, which returned to control levels, were responsible for the lack of a sustained protective effect in CCR2-deficient dystrophic mice. Similar to Ly6c^+^ bone marrow monocytes, SkMRMs that were adoptively transferred into dystrophic muscle differentiated into gal-3^+^ macrophages, an activation state with putative fibrogenic activity. We propose that the sustained differentiation of dystrophic SkMRMs cooperates with the continuous recruitment of monocytes to promote fibrosis throughout the course of muscular dystrophy. In the absence of Ly6c^hi^ inflammatory monocyte recruitment, SkMRMs and Ly6c^lo^ monocytes, become the dominant populations that promote fibrosis by adopting the gal-3^+^ program. Our observations that the transcriptional state of SkMRMs is associated with the ECM (fig. S5) and this program is retained in gal-3^+^ macrophages (fig. 3L), suggest that SkMRMs are poised to differentiate into a population with an intrinsic quality to promote fibrosis. Temporal studies will be required to determine whether recruited monocytes adopt a similar pro-fibrotic program with disease progression, as well as the development of lineage tracing systems to parse out the contribution of SkMRMs and MDMs to the gal-3^+^ macrophage pool and muscle fibrosis.

Collectively, this study identified diverse subsets of muscle macrophages with distinct functions and transcriptional profiles. The gal-3^+^spp1^+^ signature reflected the predominant transcriptional state of a dystrophic muscle macrophage. Colocalization of gal-3^+^ macrophages with stromal progenitors, and the observation that Spp1 mediates communication between these cell types, reinforces the importance of a macrophage-FAP fibrogenic axis in promoting the pathogenesis of muscle disease. This view is further supported by the observation that a similar fibrogenic axis exists in human, as gal-3^+^ macrophages were elevated in several human muscle diseases. Further, gal-3^+^ macrophages were identified in three different models of chronic muscle disease and acute muscle injury, suggesting that a canonical mechanism associated with muscle damage triggers differentiation into the gal-3^+^ state. A likely candidate is phagocytosis of apoptotic and necrotic cell debris, which has been documented as a key contributor to macrophage activation (Arnold et al., 2007). However, this represents only one of the many macrophage-FAP fibrogenic circuits that have been previously documented in muscle (Juban et al., 2018; Lemos et al., 2015). Further studies are necessary to test the effect of phagocytosis on the induction of the gal-3^+^ phenotype in diseased muscle. Mechanistic approaches relying on mouse genetics to study macrophage and stromal progenitor interactions will also advance our understanding of how this axis promotes muscle fibrosis during muscular dystrophy. In summary, by defining the transcriptional heterogeneity of muscle macrophages, this study has advanced the understanding of macrophage activation and function during muscle homeostasis and degenerative disease. The defined transcriptional states open a path for develop novel therapeutic approaches to inhibit immune-mediated muscle fibrosis.

## MATERIALS AND METHODS

### Study design

This study aimed to determine how skeletal muscle macrophages promote fibrosis by using transcriptomics to define their homeostatic signature and how this state is altered with muscle disease. Single-cell RNAseq was used to define macrophage diversity in healthy and diseased muscle. Bulk RNAseq analysis was performed on the predominant macrophage populations identified in the scRNAseq studies, to define the regulation of the transition from homeostatic to the diseased state. Spatial transcriptomics was used to understand how dystrophic macrophages interfaced with the dystrophic environment to promote fibrosis. Multiple mouse models of muscle disease were used to demonstrate the significance of a novel gal-3^+^ macrophage population in muscular dystrophy. Further, adoptive transfer experiments were used to determine if resident macrophages and peripheral monocytes both gave rise to gal-3^+^ macrophages. The translation of these studies was assessed by quantifying gal-3^+^ macrophages in human muscle disease through an immunohistochemical examination of archived, de-identified muscle biopsies. Prior approval for collecting muscle tissue and its use in research was given by the Institutional Review Board at the University of California Irvine (HS 2019-5134). All participants provided written informed consent and Health Insurance Portability and Accountability Act authorization for data collection and the use of muscle tissue for research.

### Experimental animals

C57BL/10 (#:000476), mdx mice (C57BL/10ScSn-Dmdmdx/J) (#:001801) and mdx mice in the DBA2/J background (D2-mdx) (#:013141) were obtained from The Jackson Laboratory. CD45.1 congenic mice (#:002014) were also obtained from The Jackson Laboratory and crossed with mdx mice at the University of California Irvine (UCI). B6A/J (#:012767) and C57BL/6 mice (#:000664) were bred in vivariums at Children’s National Hospital. VCP (*53*) and DUX4 (*54*) mice were provided by collaborators at UCI and Ohio State University, respectively. Animal experiments were approved by the Institutional Animal Care and Use Committee of UCI.

### Statistical analyses

Data were expressed as Mean ± SEM. Statistical analyses were performed using Graphpad Prism version 9.2. Statistical comparisons were performed using an unpaired two-tailed Student t-test. One-way or two-way ANOVA with a post-hoc Bonferroni test was conducted when comparing multiple groups. P values ≤ 0.05 were considered significant.

## List of Supplementary Materials

**Supplementary Materials and Methods**

**Supplementary Figures**

Fig. S1. Unsupervised learning and clustering of muscle macrophage scRNAseq data.

Fig. S2. Cluster 2 macrophages in dystrophic muscle resemble monocytes.

Fig. S3. Characterization of SkMRM, gal-3+ macrophages and MDMs.

Fig. S4. Prevalence of SkMRM, MDM and gal-3+ macrophages in healthy tissues.

Fig. S5. Skeletal muscle resident macrophages express a transcriptome associated with muscle homeostasis and function.

Fig. S6. Novel skeletal muscle macrophages populations heterogeneously display qualities of M1 and M2 macrophages.

Fig. S7. Top 5 and regeneration/repair-associated GO terms.

Fig. S8. Expression of Spp1 and its receptors in WT and mdx FAPs and macrophages.

Fig. S9. Regulation and localization of SkMRM and galectin-3+ macrophages in muscular dystrophy

Fig. S10. Reference-based integration of muscle macrophages scRNAseq datasets.

Fig. S11. Histological examination of acutely-injured muscle.

Fig. S12. Preferential expression of clusters 0, 1 and 2 macrophage marker genes.

## Supplementary Table

Table S1. Bulk RNASeq Summary Table

## Acknowledgments

We thank the UCI Institute for Immunology flow cytometry core for access to equipment and the UCI Institute for Clinical and Translational Science for resources that supported this study. Part of the RNAseq studies was made possible through access to the Genomics High Throughput Facility Shared Resource of the Cancer Center Support Grant (P30CA-062203) at UCI

## Funding

Research reported in this publication was supported by the: National Institutes of Neurological Disorders and Stroke (NINDS) grant R01NS120060 and National Center for Advancing Translational Sciences (NCATS) grant KL2TR001416 (to S.A.V.)

National Institute of General Medical Sciences (NIGMS) grant R01GM143536 (to J.R.Z.) National Institute of Arthritis and Musculoskeletal and Skin Diseases (NIAMS) grant U54 AR052646-07 (to M.S.).

National Institute of Arthritis and Musculoskeletal and Skin Diseases (NIAMS) grant R01AR078340 (to T.M.)

Department of Defense (DoD) Congressionally Directed Medical Research Program (CDMRP) Discovery Award W81XWH1910012 (to G.C.)

National Institute of Allergy and Infectious Diseases (NIAID) grant 1R01AI168063-01 (to S.O.) NIAMS grant P30AR075047, National Science Foundation (NSF) grant DMS11763272 and a Simons Foundation grant 594598 (to Q.N.)

## Author contributions

Conceptualization: GC, JKM, JRZ, SAV

Methodology: GC, JKM, DJ, QN, NP, QS, TM, MS, JRZ, SAV

Investigation: GC, DJ, CGJ, JMK, PKF, QN, NP, JD, AS, EM, MD, SO, ALM, QN, IM, VK, SH, TM, MWH, SB, JKJ, KK, QS, MS, MJS, JRZ, SAV

Visualization: GC, JKM, MS, JRZ, SAV

Funding acquisition: TM, MJS, JRZ, SAV

Project administration: SAV

Supervision: JRZ, SAV

Data curation: JKM, CGJ, JRZ, QS

Statistical and other formal analysis: GC, JKM, CGJ, ALM, QS, JRZ

Writing – original draft: GC, MS, JRZ, SAV

Writing – review & editing: GC, DJ, PFK, TM, MS, MJS, JRZ, SAV

## Competing interests

Dr. Mozaffar discloses an advisory role for and/or receiving research funds from Alexion, Amicus, Argenx, Arvinas, Audentes, AvroBio, Horizon Therapeutics, Immunovant, Maze Therapeutics, Momenta (now Janssen), Sanofi-Genzyme, Sarepta, Spark Therapeutics, UCB, and Modis/Zogenix. Dr. Mozaffar also serves on the data safety monitoring board for Acceleron, Avexis, and Sarepta. Drs. Golann, Si and Stec are employees and shareholders of Regeneron Pharmaceuticals. Dr. Spencer is a co-founder of MyoGene Bio and SkyGene Bio. All other authors declare that they have no competing interests.

## Data and materials availability

The RNA sequencing data will be deposited in the Gene Expression Omnibus (GOE) managed by the National Center for Biotechnology Information.

