## Supplementary materials for "Single-cell and spatial transcriptomics identify a macrophage population associated with skeletal muscle fibrosis"

### List of Supplementary Materials

**Supplementary Materials and Methods**

**Supplementary Figures**

Fig. S1. Unsupervised learning and clustering of muscle macrophage scRNAseq data.

Fig. S2. Cluster 2 macrophages in dystrophic muscle resemble monocytes.

Fig. S3. Characterization of SkMRM, gal-3+ macrophages and MDMs.

Fig. S4. Prevalence of SkMRM, MDM and gal-3+ macrophages in healthy tissues.

Fig. S5. Skeletal muscle resident macrophages express a transcriptome associated with muscle homeostasis and function.

Fig. S6. Novel skeletal muscle macrophages populations heterogeneously display qualities of M1 and M2 macrophages.

Fig. S7. Top 5 and regeneration/repair-associated GO terms.

Fig. S8. Expression of Spp1 and its receptors in WT and mdx FAPs and macrophages.

Fig. S9. Regulation and localization of SkMRM and galectin-3+ macrophages in muscular dystrophy

Fig. S10. Reference-based integration of muscle macrophages scRNAseq datasets.

Fig. S11. Histological examination of acutely-injured muscle.

Fig. S12. Preferential expression of clusters 0, 1 and 2 macrophage marker genes.

**Supplementary Table**

Table S1. Bulk RNASeq Summary Table

### SUPPLEMENTARY MATERIALS and METHODS

#### Single-cell RNA sequencing

*Single-cell preparation and analysis of muscle macrophages.*

Sorted cells were washed and resuspended at a concentration of ~1000 cells/μL. Using the 10x genomics platform, libraries for WT and mdx muscle macrophages were generated following the Chromium Single Cell 3′ Reagents Kits v2 User Guide: CG00052 Rev B. Quantification of cDNA libraries was performed using Qubit dsDNA HS Assay Kit (Life Technologies Q32851) and high-sensitivity DNA chips (Agilent 5067-4626). Quantification of library construction was performed using KAPA qPCR (Kapa Biosystems KK4824). 10x libraries were sequenced on the Illumina HiSeq4000 platform to achieve an average of ~50,000 reads per cell according to recommendations in the Chromium Single Cell 3′ Reagents Kits v2 User Guide: CG00052 Rev B. Sequencing reads were processed utilizing the 10x Genomics Cell Ranger 2.1.0. Each library was aligned to an indexed mm10 genome using Cell Ranger Count. To generate an aggregated matrix of WT and mdx cells and to prepare data for downstream analysis, the Cell Ranger Aggr function was used to normalize the number of confidently mapped reads per cell each library.

To identify cell clusters, the Seurat pipeline (Version 3.0.2) was applied to the aligned cell matrix using R (Version 3.6.1). Quality control filtering was first performed to remove genes that were not expressed (>0) in at least three cells and cells that had less than 200 genes. The trimmed expression count matrix was log-transformed for downstream processing and highly variable genes were detected. Principal Component Analysis was combined with the elbow method to determine loadings to be used for the generation of Uniform Manifold Approximation and Projections (UMAP) with 10 Principal Components (PCs) included. Using these same PCs, Seurat's default clustering was performed and was followed up with marker gene detection to elucidate gene expression signatures corresponding to the resultant clusters.

*Reference-based mapping of macrophages, FAP, and tenocytes*

We performed anchoring and integration of single-cell datasets of uninjured WT (i.e. Day 0) (*1*), and mdx mouse muscle that served as a negative control group in a separate study of mice treated with tamoxifen (dataset provided by the MJ Spencer). Seurat (Version 3.2.2, R Studio version 3.6.1) was used for anchoring and integration (*2*). Briefly, merged Seurat objects were normalized, and highly variable genes (features) and scaling were performed with SCTransform (*3*). The top 2,000 highly variable features were selected and used for anchoring. Integration anchors (30 dimensions) were computed and used for integration. For neighbor and cluster identification, the integrated object was scaled, and significant principal components (PCs) were identified via statistical and heuristic testing as recommended in Seurat. Clustered cells were visualized using UMAP. Prior to anchoring and integration, macrophage identities from both datasets were renamed to match macrophage identities from the current study. Briefly, cells from WT (Monocyte_patrolling, monocyte_inflammatory, monocyte_mixed, M2 macrophage_*Cx3cr1*$^{lo}$, M2 macrophage_*Cx3cr*$^{hi}$, and dendritic cells) and mdx (*Lyz2*, *Ctss*, *Cd68*, *Fcgr2b*, *Cd14*, and *Adgre1*-expressing cells) were subclustered and subjected to reference-based mapping using the macrophage identities described in this study (i.e., reference cells – Macrophage 0-5) (*4*). Following a similar approach, WT FAP_adipogenic, FAP_pro-remodeling, FAP_stem, and tenocytes (*1*) were used as reference to assign identities to *Pdgfra*$^{+}$ FAPs and tenocytes in the mdx dataset. All other previously assigned cell identities described in Oprescu *et al*. (*1*) were kept in the final clustering with the exception of capillary, mixed, and vein endothelial cells, which were collapsed into “endothelial cells”; T & NK cells, which were separated into T and NKT cells; and cycling cells, which were identified by expression of *Mki67*.

*Reference-based mapping of B6AJ and Limb Girdle macrophages*

We performed reference-based mapping, anchoring and integration of 8-month-old B6A/J (LGMD2B, n=2) and B6 (WT, n=1) muscle datasets using Seurat (Version 3.2.2, R Studio version 3.6.1) (*4*). The annotated macrophage dataset in Fig. 1 was used as the reference. WT B10, mdx, WT B6 and B6A/J objects were merged, SCTransform-normalized, anchored, and integrated as described above. Clustered cells were visualized using two-dimensional UMAP.

*Modeling cell-cell communication networks*

Intra- and intercellular communication networks were modeled based on the abundance of known ligand-receptor transcript pairs with CellChat (version 1.1.3) (*5*). To identify conserved and perturbed FAP-macrophage communication networks in WT and mdx muscles, we lifted cells from a WT-mdx integrated object and performed joint manifold and classification learning analyses as described in CellChat.

**Bulk RNA sequencing and analysis**

Total RNA was monitored for quality control using the Agilent Bioanalyzer Pico RNA chip (Agilent Technologies, Santa Clara CA) and Nanodrop (Thermo Fisher Scientific, Waltham MA) absorbance ratios for 260/280nm and 260/230nm. Library construction was performed according to the SMARTer Stranded Total RNA-Seq Kit v2- Pico Input Mammalian (Takara Bio, Mountain View CA). The input quantity for total RNA was 2ng. Total RNA was fragmented for 3min at 94°C. SMART (Switching Mechanism At 5’ end of RNA Template) cDNA synthesis technology was used to synthesize cDNA from the fragmented total RNA. Illumina adapters were ligated to the ends, and enriched by 5 cycles of PCR. R-probes v2 (mammalian-specific) are then hybridized to the cDNA that contains ribosomal RNA and human mitochondrial rRNA sequences. The R-probe v2 hybridized cDNA are then cut by ZapR v2. The leftover library fragments are further enriched by 14 cycles of PCR. The final libraries are purified via AMPure XP beads. The resulting libraries were validated by qPCR (Kapa library quantification kit, Kapa Biosystems, Willimington MA) and sized by Agilent Bioanalyzer DNA high sensitivity chip. The concentrations for the libraries were normalized and then multiplexed together. The multiplexed libraries were sequenced using paired-end 100 chemistry on the NovaSeq 6000 (Illumina, San Diego CA).

Public data for Microglia (*6*) (GSE132877) and bone-derived macrophage (*7*) (provided by authors) RNA-seq were processed from original fastq read files. All reads were mapped to the mouse genome (mm10) (*8*) with TopHat (*9*) (version 2.0.14) with reference GENCODE transcript annotation (*10*) (M9). Overlapping reads were counted and summarized by gene using HTSeq (*11*) (1.99.2). To determine DEGs between datasets, the R package DESeq2 (*12*) (version 1.28.1) was used. From DESeq2 output, DEGs were classified for 2-fold changes up or down with a fdr < 0.01. Gene sets from the Molecular Signature Database (*13*) (mSigDB) were downloaded from the GSEA (*14*) webpage (http://software.broadinstitute.org/gsea) to determine gene set enrichment. Plots were generated in R using VennDiagram (version 1.6.20), ggrepel (version 0.9.0), ComplexHeatmap (version 2.4.3), plyr (version 1.8.6) and ggplot2 (version 3.3.3).

**Spatial transcriptomics**

The 10x Genomics Visium Spatial Gene Expression platform was used for spatial transcriptomics analysis of muscle from dystrophic mice (D2-mdx, stock #013141 from the Jackson Laboratories) according to manufacturer's guidelines. In brief, 6-week-old male mice were euthanized via cervical dislocation under isoflurane anesthesia, and the gastrocnemius/plantaris muscle complex was immediately dissected and frozen in OCT embedding media in liquid nitrogen-cooled isopentane. Muscle tissue was cryosectioned at -20°C at 10µm thickness onto Visium Spatial Gene Expression slides (10x Genomics) and stored at -80°C until processing. Sections were fixed in pre-chilled methanol for 30 minutes. H&E staining was performed per the published protocol from 10x Genomics, with imaging performed using a Zeiss Axio Observer microscope. H&E images were stitched and processed using Zen 2.0 software. Following imaging, tissues were permeabilized for 12 minutes, which was predetermined as the optimal time for 10 µm mouse muscle sections using the 10x Genomics Visium Tissue Optimization Kit. Spatially-tagged cDNA libraries were built using the 10x Genomics Visium Spatial Gene Expression Library Construction Kit. Sequencing was performed on an Illumina NextSeq 500/550 using 150-cycle High Output kits (Read 1 = 28, Read 2 = 120, Index 1 = 10, and Index 2 = 10). Alignment to the mouse reference genome mm10 (Ensembl 93) was done using the Space Ranger 1.0.0 pipeline to derive a feature spot-barcode expression matrix (10x Genomics). Alignment of H&E images was done using Loupe Browser.

**RNAscope**

6-week-old male mice were euthanized via cervical dislocation under isoflurane anesthesia, and the gastrocnemius/plantaris muscle complex was immediately dissected and frozen in OCT embedding media in liquid nitrogen-cooled isopentane. Muscle tissue was cryosectioned at -20°C at 10µm thickness onto Superfrost Plus micro slides (VWR) and stored at -80°C until processing. Slides were prepared for RNAscope® assay according to Advanced Cell Diagnostics (ACD) protocol for Fresh Frozen Tissue (Document number 320513, ACD). RNAscope® was performed according to manufacturer's protocol RNAscope® Multiplex Fluorescent Assay V2 (Document

Number 320293-USM, ACD) using RNAscope® Probe - Mm-Spp1-C2 (ACD, catalog #435191-C2) with probe diluent (ACD, catalog #300041) and Amp4 Alt C-FL (Atto 550). Prior to counterstaining, the section was incubated in blocking solution (20% goat serum, 0.3% Triton X-100 in 1X PBS) with 1:100 Laminin (Sigma, L9393) for 30 minutes. Slides were washed with PBS two times for 1 minute and incubated in a filtered blocking solution with 1:250 Alexa Fluorophore 488 (ThermoFisher, 32731). Slides were washed with PBS one time for 1 minute then manufacturer's protocol was followed for counterstaining and then mounted using Fluoromount G (ThermoFisher, 00-4958-02). Sections were imaged using Zeiss Axioscan microscope and processed using HALO software (Indica Labs).

**Tissue cell isolation**

*Skeletal muscle*

Single cell isolation from hind-limb muscles was performed as previously described (*15*). Briefly, following whole body perfusion with PBS, hind-limb muscles were harvested from the mice while removing the popliteal lymph node to exclude any potential contamination of non-muscle-residing lymphocytes in our preparations. Wild type mdx hind limb musculatures of each mouse were processed and analyzed individually unless specified otherwise. Minced hind limb muscles were submitted for 2 x 30-minute rounds of enzymatic digestion with 0.2 mg/ml collagenase P (Roche) and 20 μg/ml of DNAse (Roche) in serum-free DMEM. Following digestion, single-cell preparations were sequentially filtered through a 70 and 40 μm filter basket. The filtrate was suspended over 5 ml of Histopaque 1077 (Stem cell solutions) and centrifuged at ~315 x g for 30 minutes at room temperature. Cells at the interface were harvested, counted, and analyzed or sorted by flow cytometry.

*Heart and quadriceps*

Immune cells from a single muscle such as heart or quadriceps were isolated using a modification of the previously described procedure. Heart or quadriceps was minced then digested 2 x 20 minutes in a 0.2 mg/ml collagenase P solution supplemented with 20 µg/ml of DNase. The muscle suspension is then sequentially filtered through a 70 and 40 µm filter-basket. Following a final suspension using a 35-µm strainer mesh, immune cells are counted and stained for flow cytometry analysis.

*Liver*

Immune cells from the liver were isolated as described (*16*) with some modifications. Briefly, following whole body perfusion with PBS, liver lobes were minced and digested in 0.2 mg/ml collagenase P (Roche) and 20 µg/ml of DNAse (Roche) in serum-free DMEM at 37 C for two rounds of 20 minutes enzymatic digestion. Red blood cells were lysed using 1X RBC lysis buffer. Cell pellets were washed with 1X PBS, resuspended and filtered through 70 and 40 µm filter-baskets. Cells were counted prior to cell surface staining.

*Bone marrow*

Primary mouse bone marrow cells were isolated by flushing the tibiae and femurs with phosphate-buffered saline (PBS). The resulting cell suspension was gently disaggregated and passed through a 70 µm cell strainer to produce a single cell suspension. Red blood cells were lysed by incubating for 5 minutes with ACK lysis buffer. Bone marrow cells were counted prior to cell surface staining.

*Mouse PBMCs*

Mouse blood was collected by cardiac puncture in an Eppendorf tube containing 10 µl of Heparin. Briefly, the blood was diluted to a 1:1 volume ratio with PBS and gently layered on top of the density gradient medium Histopaque 1077 (Stem cell solutions). Cells were harvested following a 20 min centrifugation at 315 x g for 20 minutes. PBMCS were then washed twice with PBS prior to downstream applications.

*Brain*

Central nervous system (CNS) macrophages were sorted from the single-cell suspension obtained separately from the brain and spinal cord as described previously (*17*). Briefly, brains and spinal cords were dissected from PBS-perfused mice and mechanically dissociated using a Dounce grinder (# K8853000007, ThermoFisher) to obtain tissue homogenate. CNS tissue homogenates were further digested with Collagenase IV (Worthington-Biochemical) (1 mg/ml) and DNase I (Sigma Aldrich) (200 Units/ml) in 5 ml of RPMI 1640 media for 45 minutes at 37°C. To separate myelin debris from cells, digested tissue was first passed through a 100 µm cell strainer then 23% Percoll (GE Life Sciences Cat# 17089101) density gradient centrifugation was used (400g without brakes for 25 min at 4° C). The myelin layer at the interface was removed and the single cells in the pellet were collected. Additional RBC lysis was performed using ACK buffer (# A1049201, ThermoFisher) for 60 seconds on ice when necessary. The single-cell suspension was then passed through a 70 µm cell strainer, washed twice with PBS for subsequent FACS and RNASeq analysis.

*Skin*

To obtain single cell suspension, minced whole skin samples from shaved mice were digested with 10 ml of a solution containing 0.25% collagenase (Sigma, C9091), 0.01 M HEPES (ThermoFisher, BP310), 0.001 M sodium pyruvate (ThermoFisher, BP356), and 0.1 mg/mL DNase (Sigma, DN25) at 37 ºC for 1 hour with rotation, and then filtered through a 70-µm and 40-µm filter, spun down, and resuspended in 2% fetal bovine serum (Alphabioregen, Alpha FBS). Isolated cells were counted prior to cell staining

*Lung*

Single-cell suspensions from the lungs of mice were isolated using mechanical and enzymatic digestion. The trachea was exposed and 1 ml of digestion media composed of RPMI containing 0.1mg/ml collagenase P and 0.02 mg/ml DNase was injected into the trachea to inflate the lungs. The lungs were then dissected, minced, and digested in 3 ml of digestion media for 45 minutes while rocking at 37°C. Following digestion, suspensions were diluted with HBSS and filtered through 70⌠m filter and pelleted using centrifugation. Red blood cells were lysed by incubating cells with ACK lysis buffer (Life Technologies) for 5 minutes. Single-cell suspensions were then analyzed using flow cytometry

**Barium Chloride injury**

Prior to muscle injury, 6-7 weeks, mice were anesthetized using a mixture of 2-3% L/min of isoflurane and oxygen. Barium Chloride-induced acute muscle injury was carried out by 4, 20 µl intramuscular injections of barium chloride dissolved in saline solution (1.2% w/v), applied along the length of the quadriceps to maximize the distribution of barium chloride. The contralateral quadriceps served as untreated control. Following injury, animals were euthanized at different time points, and quadriceps were harvested for flow cytometry or histological analyses.

**Adoptive transfer**

Monocytes were isolated from bone marrow using EasySep Mouse Monocyte Enrichment kit (Stem Cell Technologies). Briefly, femurs and tibias were harvested and flushed through with cold PBS. Bone marrow cell suspensions were passed through a 70-µm cell strainer to obtain a single cell suspension. Red blood cells were removed using the RBC lysis buffer according to the manufacturer's instructions (Sigma-Aldrich). The bone marrow cell suspension was treated with the EasySep reagents and monocytes isolated by depletion using an EasyPlate magnet (Stem Cell

Technologies). Muscle immune cells were isolated from hind limb muscles as described previously and enriched using the EasySep Release Mouse APC selection kit (Stem Cell Technologies). Briefly, the muscle single cell suspension was resuspended in 0.25 mL recommended medium and incubated with the Folr2-APC antibody (2 µg/ml) for 5 minutes on ice. Following an incubation with 25 µl of APC selection cocktail for 5 minutes at 4 C, APC+ cells selection is obtained by incubating with RapidSpheres magnetic beads on ice for 3 minutes. $Folr2^+$ macrophages bound to the magnetic beads were eluted after the addition of the ice-cold release buffer on ice for 3 minutes, counted and saved for downstream experiments. Isolated monocytes (20000 cells/µl in PBS) or $Folr2^+$ macrophages (15000 cells/µl in PBS) were injected in 10 µl intramuscularly. Briefly, mdx mice were anesthetized with isoflurane while placed over a heating pad to maintain thermoregulation. Anesthetized mice were sterilized with 70% ethanol and 10 µl of cell suspension were injected into the quadriceps using a Hamilton syringe. When appropriate, the other quadriceps was used as a control where no cells were injected into the muscle. Flow analysis of the quadriceps of mdx mice was performed 2 and 7 days following the adoptive transfer.

**Histological analyses**

*Sirius Red*

Muscle cryosections were fixed overnight in Bouin's fluid at room temperature and cleared in xylene prior to hydration with decreasing ethanol concentrations. Sections were stained for 15 minutes at room temperature under agitation in 0.02% Sirius red solution. Following washes in acetic acid, the slides are dehydrated and mounted with a non-aqueous mounting media. The collagen content labeled by the Picrosirius red staining is examined using a Cytation 5 brightfield microscope.

*Hematoxylin and Eosin*

Hematoxylin and Eosin stain was performed on fresh frozen muscle tissues. Quadriceps muscle cryosections (8 µm) were stained with Harris hematoxylin and then counterstained with eosin. Sections were then examined using a Cytation 5 brightfield microscope.

*Immunofluorescence*

For IHC images, serial sections were first fixed with 4% paraformaldehyde for 15 min. Slides were washed with PBS three times for 5 minutes and incubated in blocking solution for 1 hour. Slides were then washed with PBS three times for 5 minutes and incubated in primary antibody solution (primary at 1:100 in blocking solution) overnight in a humid box. Primary antibodies were as follows: Laminin (Sigma-Aldrich, L9393 or Abcam, ab11576 at 1:500), Collagen I (Abcam, ab270993), F4/80 (Abcam, ab6640). Slides were then washed three times for 5 minutes with PBS and then incubated in filtered secondary antibody solution (secondary at 1:250 in blocking solution). Secondary antibodies were as follows: rabbit 546 (ThermoFisher, 11035), rat 488 (ThermoFisher, 11006), rat 647 (ThermoFisher, 21247). Slides were washed once with PBS for 5 minutes and then incubated in Hoechst 33342 (ThermoFisher, H3570) for 5 minutes. After two additional 5 minutes PBS washes, slides were mounted in Fluoromount G (ThermoFisher, 00-4958-02), and cover slipped. Sections were imaged using Zeiss Axio Scan microscope and processed using HALO software.

**RNA isolation and qPCR analysis**

Liquid nitrogen-frozen muscle samples were homogenized, and RNA was extracted using TRIsure (Bioline) and the Quick-RNA Miniprep kit (Zymo Research) per manufacturer instructions. Cell populations were sorted directly into lysis buffer, and RNA was isolated using the Quick-RNA Microprep kit (Zymo Research). Complementary DNA (cDNA) was synthesized from 150 ng (sorted cells) or 1000 ng (whole muscle) of DNase-treated RNA using the SensiFAST cDNA synthesis kit (Bioline). Gene expression was quantified using TaqMan expression assay probes (Life Technologies) and 2x SensiFAST probe No-ROX mix (Bioline). All gene expression was normalized to 18s unless otherwise noted.

**Flow cytometry**

To discriminate between live and dead cells, cells are resuspended in 100 µl of Zombie NIR viability dye (1:1000 in 1X PBS) for 30 minutes on ice while protected from light. Fc receptor blocking of muscle single-cell suspensions was performed by incubating cells with an anti-CD16/32 antibody (clone 2.4G2) prior to staining. Single-cell suspensions were stained with a panel of antibodies against several cell surface antigens (see table #?) Analysis was performed on live cells on a BD FACSAria Fusion flow cytometer with FACSDiva software (BD Bioscience). Post-acquisition analysis was performed with Flowjo software version 10.8.

### SUPPLEMENTARY FIGURES

#### Supplementary Figure 1

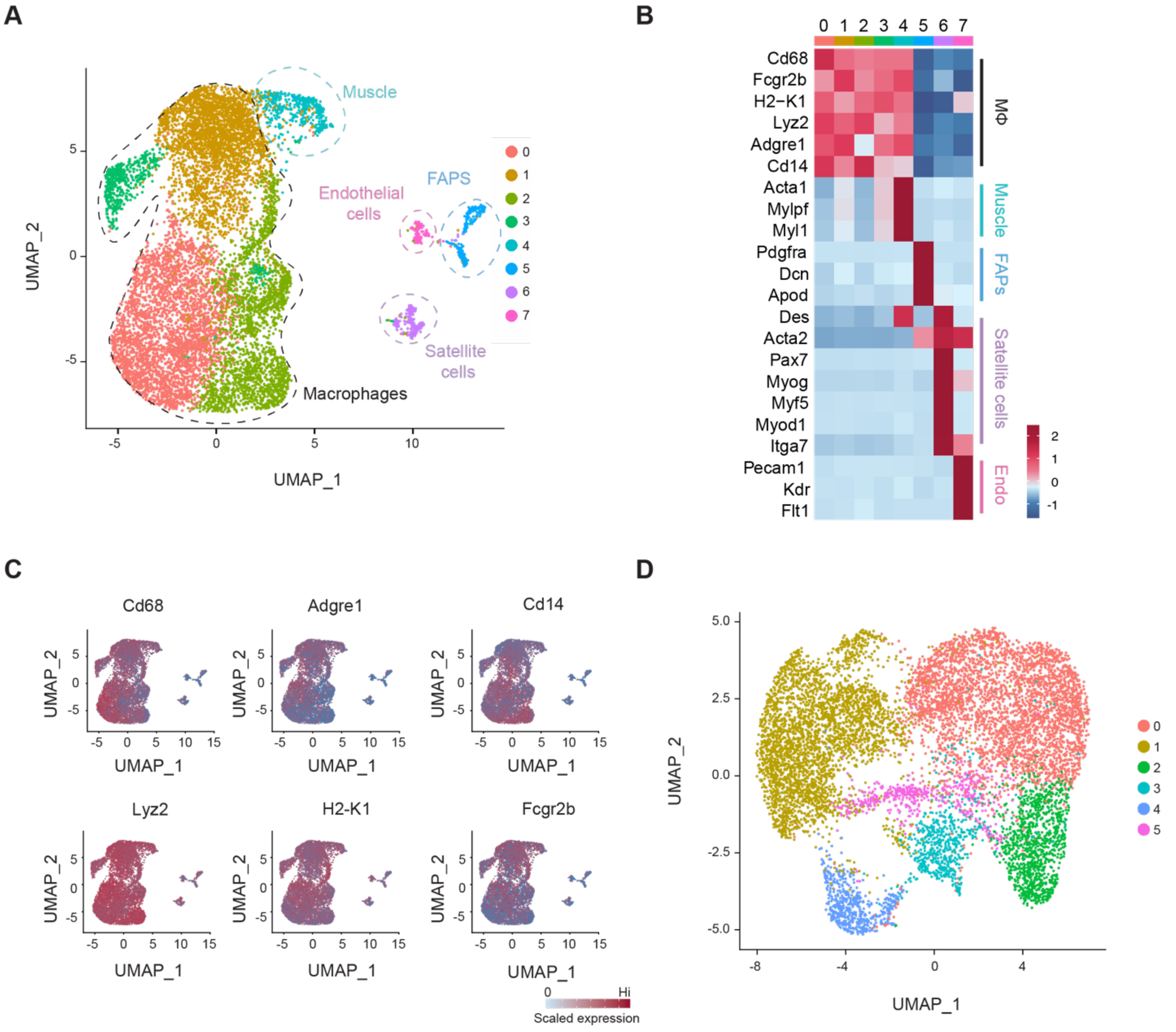


**Fig. S1. Unsupervised learning and clustering of muscle macrophage scRNAseq data.** (**A**) Dimensionality reduction via UMAP showing all cells in WT and mdx muscle macrophage datasets. (**B** and **C**) heatmap showing cell-specific marker genes (B) and feature plots showing the expression of macrophage marker genes (C). (**D**) UMAP showing sub-clustering analysis of bulk macrophages. Six novel populations were identified.

### Supplementary Figure 2

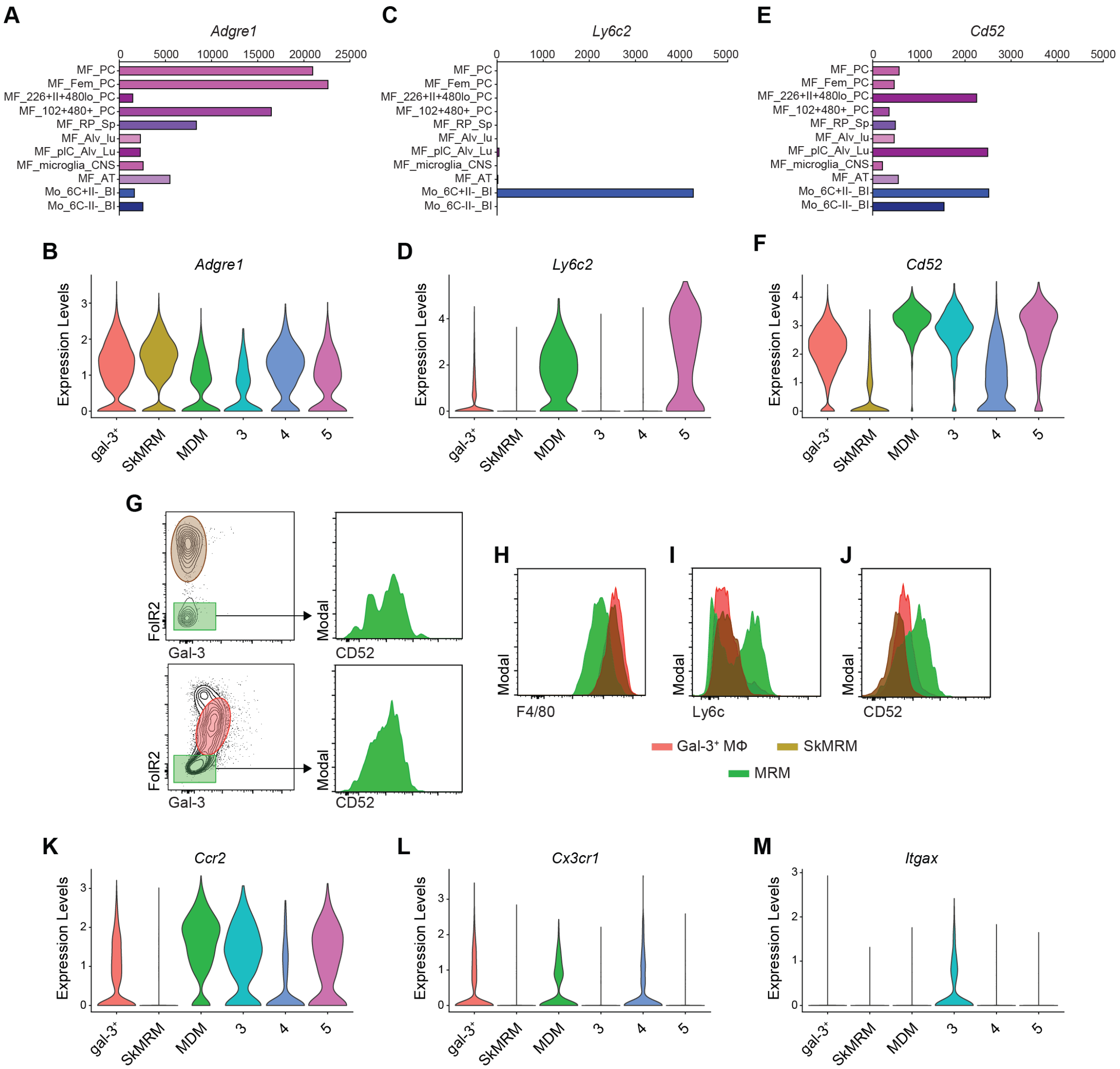


**Fig. S2. Cluster 2 macrophages in dystrophic muscle resemble monocytes.** (**A-F**) Expression of key monocytes markers. The expression of levels of *Adgre1* (F4/80) (A), *Ly6c* (C) and *CD52* (D) in monocyte and macrophages populations were assessed through the ImmGen database (www.immgen.org). Violin plots showing expression of *Adgre1* (B), *Ly6c* (D) and *CD52* (F) in the scRNAseq analysis (Fig. 1). MF.Fem.PC= Female Peritoneal Macrophages obtained by sorting F4/80$^+$ICAM$^+$CD5$^-$CD19$^-$CD43$^-$ cells; MF.226+II+480lo.PC= Peritoneal Small Macrophages obtained by sorting CD115$^+$CD11b$^+$F4/80$^{lo}$CD102$^{lo}$MHCII$^+$CD226$^+$; MF.102+480+.PC= Peritoneal Large Macrophages obtained by sorting

$CD115^+CD11b^+F4/80^+CD102^+MHCII^{lo}CD226^-$ ; MF.RP.Sp= Red pulp Macrophages from spleen sorted on $CD11b^{lo}F4/80^+Mertk^+CD64^+$; MF.Alv.Lu= Lung Alveolar Macrophages sorted on $CD45^+CD11c^+SiglecF^+$; MF.pIC.Alv.Lu= Lung Alveolar Macrophages stimulated with Polyinosinic-polycytidylic acid [poly(I:C)]; MF.microglia.CNS= Brain Microglia Macrophages; MF.AT= Inguinal and Perigonadal Adipose Tissue Macrophages; Mo.6C+II-.Bl= $Ly6C^{hi}$ blood monocytes sorted on $B220^-CD3^-Ly6G^-CD45^+CD115^+CD11b^+Ly6C^+$; Mo.6C-II-.Bl= $Ly6C^{lo}$ blood monocytes sorted on $B220^-CD3^-Ly6G^-CD45^+CD115^+CD11b^+Ly6C^-$. (**G-J**) Representative gating strategy for interrogating the expression of F4/80 (H), Ly6c (I) and CD52 (J) in $Gal\text{-}3^+$ Mϕ, SkMRMs and MDMs. (**K-M**) Violin plots showing the expression levels of monocyte-related genes, *Ccr2* (K), *Cx3cr1* (L) *Itgax* (CD11c) (M).

### Supplementary Figure 3

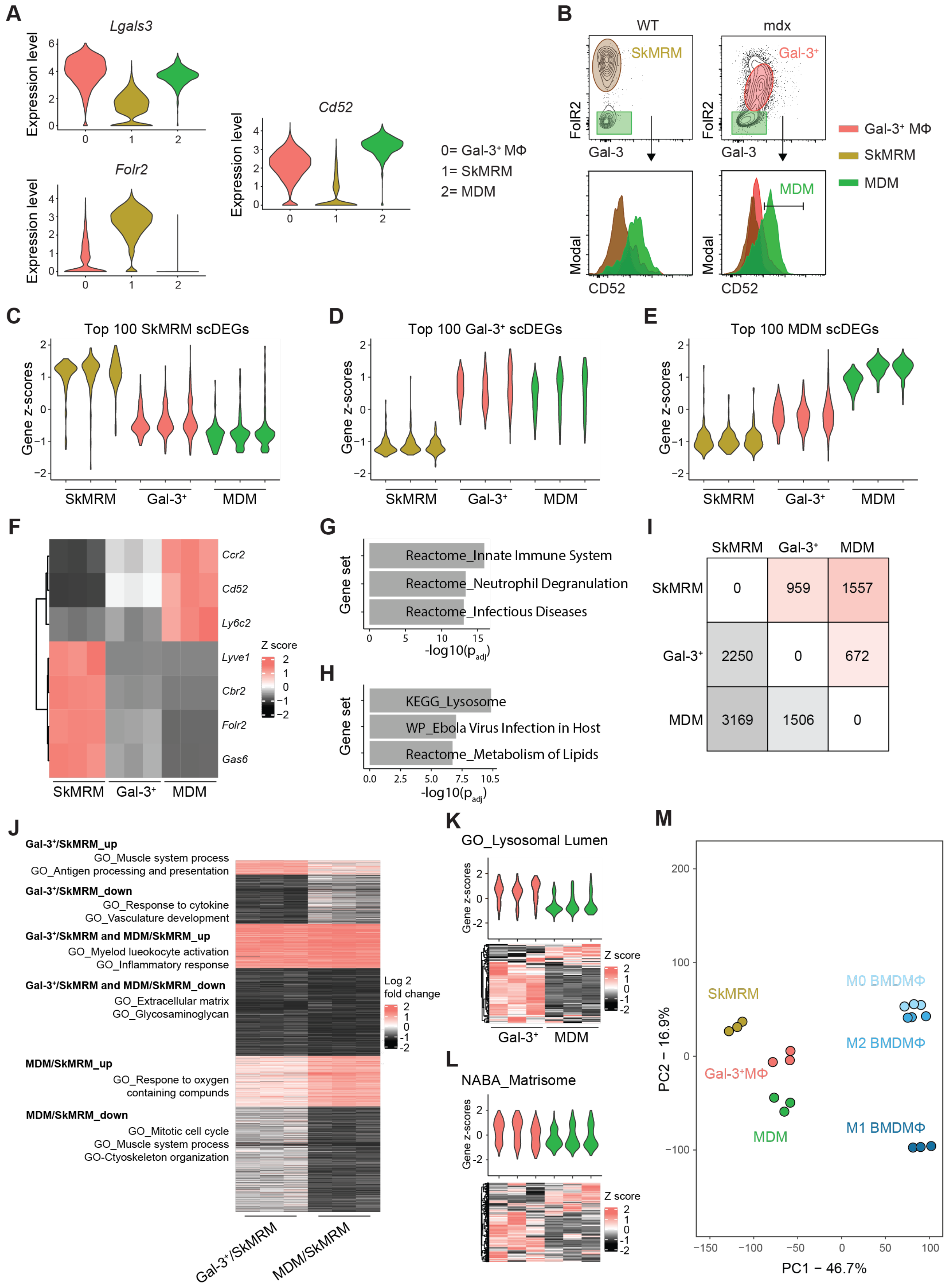


| | SkMRM | Gal-3⁺ | MDM |
|---|---|---|---|
| SkMRM | 0 | 959 | 1557 |
| Gal-3⁺ | 2250 | 0 | 672 |
| MDM | 3169 | 1506 | 0 |

**Fig. S3. Characterization of SkMRM, gal-3$^+$ macrophages and MDMs.**
**(A)** Violin plots of clusters 0 (Gal-3$^+$ macrophages), 1 (SkMRM) and 2 (MDM) marker genes. (**B**) Representative flow cytometry plots showing the gating strategy for Gal-3$^+$ Mϕ, SkMRM and MDM. (**C-E**) Violin plots showing the z-scores for the top 100 scDEGs from SkMRMs (C), Gal-3$^+$ Mϕ (D) and MDMs (E) in the transcriptomes of sorted populations. (**F**) Heatmap showing the expression of representative marker genes from the scRNAseq analysis in the transcriptomes of sorted macrophage populations. (**G and H**) Pathway analysis showing the top 3 gene sets enriched in principal component 1 (G) and principal component 2 (H) in the PCA shown in figure 2D. (**I**) A matrix showing the number of genes upregulated (pink) or downregulated (grey) in SkMRM, gal-3$^+$ Mϕ or MDMs. FDR < 0.01, FC > 2. (**J**) Heatmap with selected GO pathways enriched in gal-3$^+$ Mϕ, MDMs or both. (**K and L**) Enrichment of genes associated with the lysosomal lumen (K) and the matrisome (L) in gal-3$^+$ Mϕ or MDMs, visualized by heatmap and violin plot. (**M**) Principal component analysis applied to muscle and bone marrow-derived macrophages (BMDMϕ).

### Supplementary Figure 4

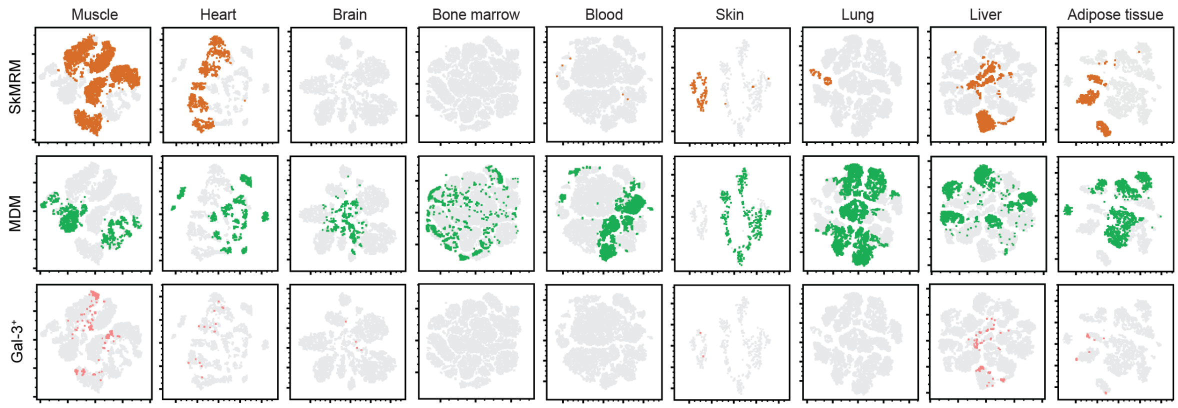


**Fig. S4. Prevalence of SkMRM, MDM and gal-3$^{+}$ macrophages in healthy tissues.** Concatenated TSNE flow plots of live CD11b$^{+}$F4/80$^{+}$Siglec-F$^{-}$ cells from 4-wk-old WT mice. n= 5 per organ or tissue. Grey shows CD11b$^{+}$F4/80$^{+}$Siglec-F$^{-}$ cells; orange indicates the proportion of CD11b$^{+}$F4/80$^{+}$Siglec-F$^{-}$ cells that are Gal-3$^{lo}$Folr2$^{hi}$; green indicates the proportion of CD11b$^{+}$F4/80$^{+}$Siglec-F$^{-}$ that are CD52$^{+}$; and pink indicates the proportion of CD11b$^{+}$F4/80$^{+}$Siglec-F$^{-}$ that are Gal-3$^{hi}$Folr2$^{lo}$.

### Supplementary Figure 5

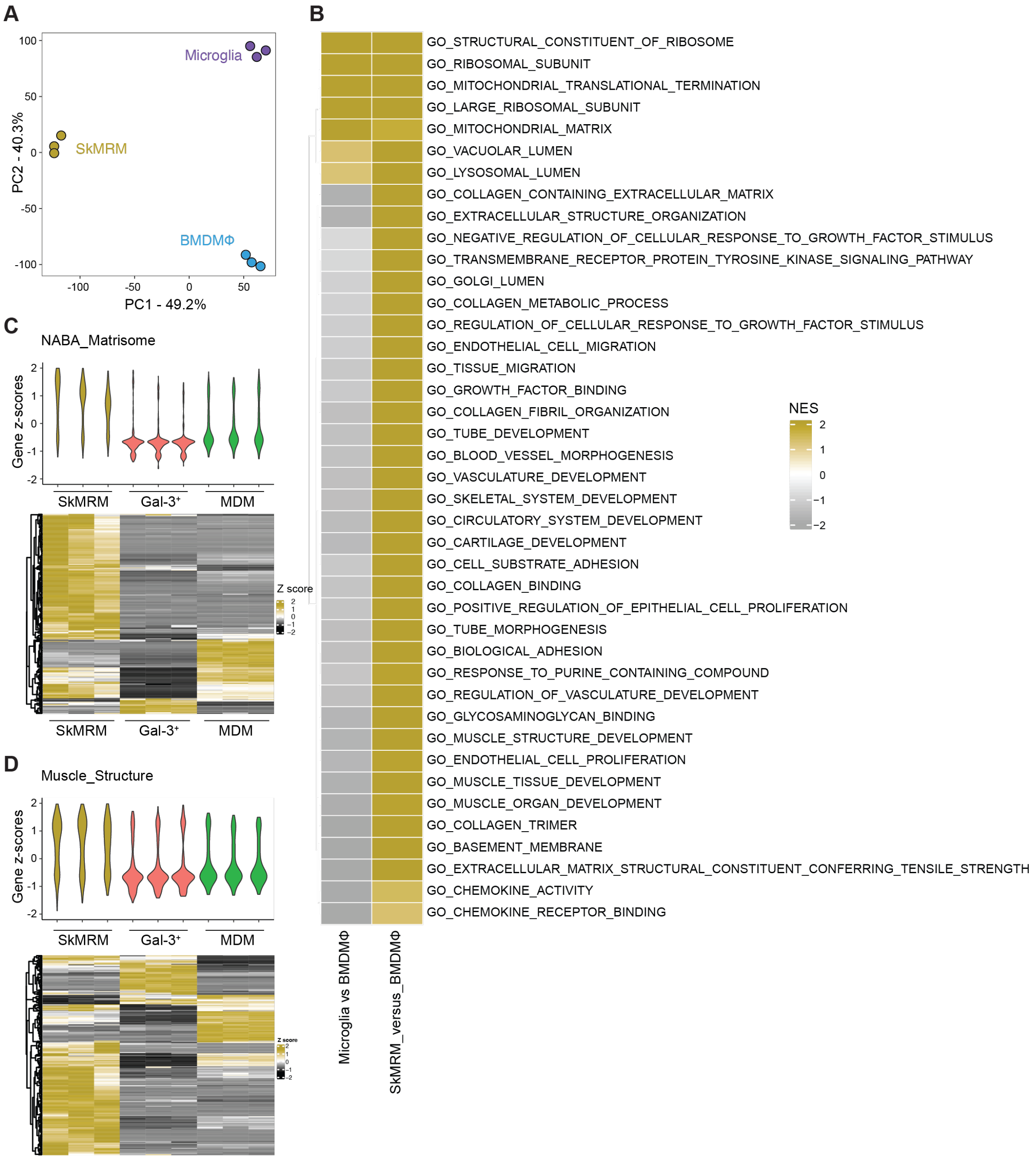


**Fig. S5. Skeletal muscle-resident macrophages express a transcriptome associated with muscle homeostasis and function.** (**A**) Principal component analysis of SkMRM, microglia and BMDMϕ. (**B**) Heatmap summarizing GSEA analysis with Gene Ontology (GO) pathways that indicate enrichment of muscle- and ECM-related pathways in SkMRMs. NES= Normalized Enrichment Score. (**C and D**) Violin plots of gene z-scores and corresponding heatmaps showing the relative expression of genes associated with the matrisome (C) and muscle structure (D) in SkMRMs.

### Supplementary Figure 6

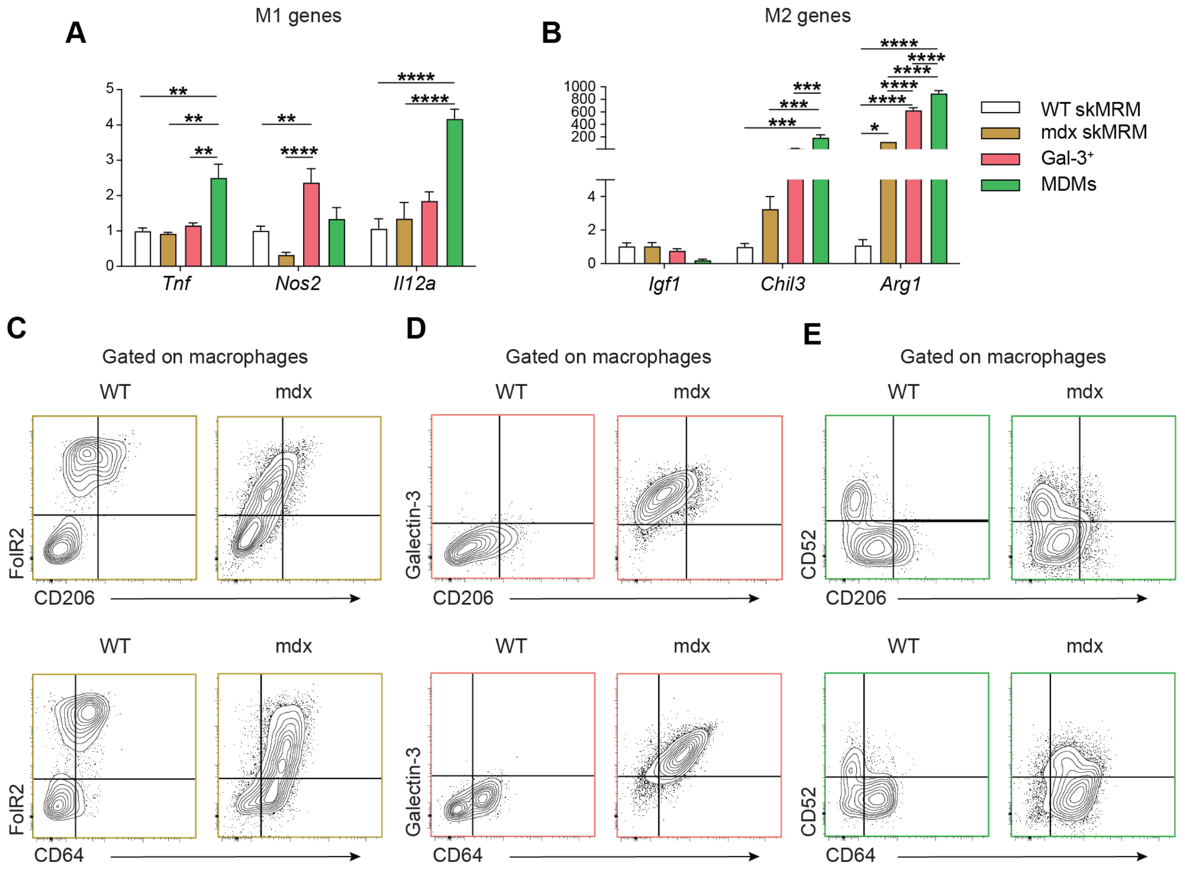


**Fig. S6. Novel skeletal muscle macrophage populations heterogeneously express qualities of M1 and M2 macrophages.** (**A** and **B**) Expression of key M1 (A) and M2 markers genes in sorted WT SkMRM, mdx SkMRM, Gal-3$^{+}$ Mϕ and MDMs from mdx mice. n= 3 sorted macrophages for each subpopulation. A two-way ANOVA with a Tukey multiple comparison test was performed. (**C-E**) Representative contour plots showing the differential expression of CD206 (M2 marker) and CD64 (M1 marker) in SkMRM, Gal-3$^{+}$ Mϕ and MDMs from 4-wk-old WT and mdx mice. n= 5. *p<0.05, **p<0.01, ***p<0.001, ****p<0.0001 using or 2-way ANOVA with Tukey's multiple comparison test (A and B).

**Supplementary Figure 7**

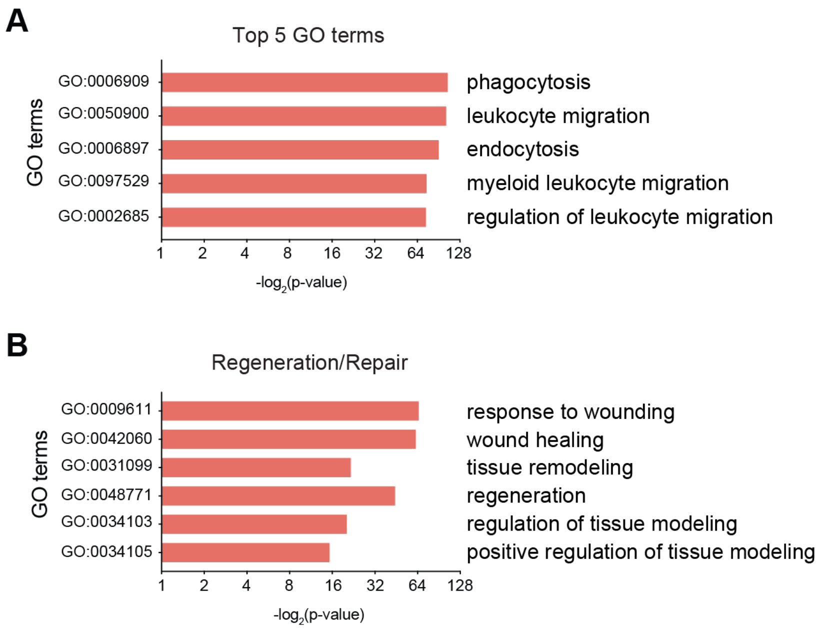


**Fig. S7. Top 5 and regeneration/repair-associated GO terms.** (**A** and **B**) Gene ontology analysis performed on the differentially expressed genes between *Lgals3*$^{lo}$ and *Lgals*$^{hi}$ spots from the spatial transcriptomics. The top 5 GO terms (A) and the regeneration and repair terms (B).

### Supplementary Figure 8

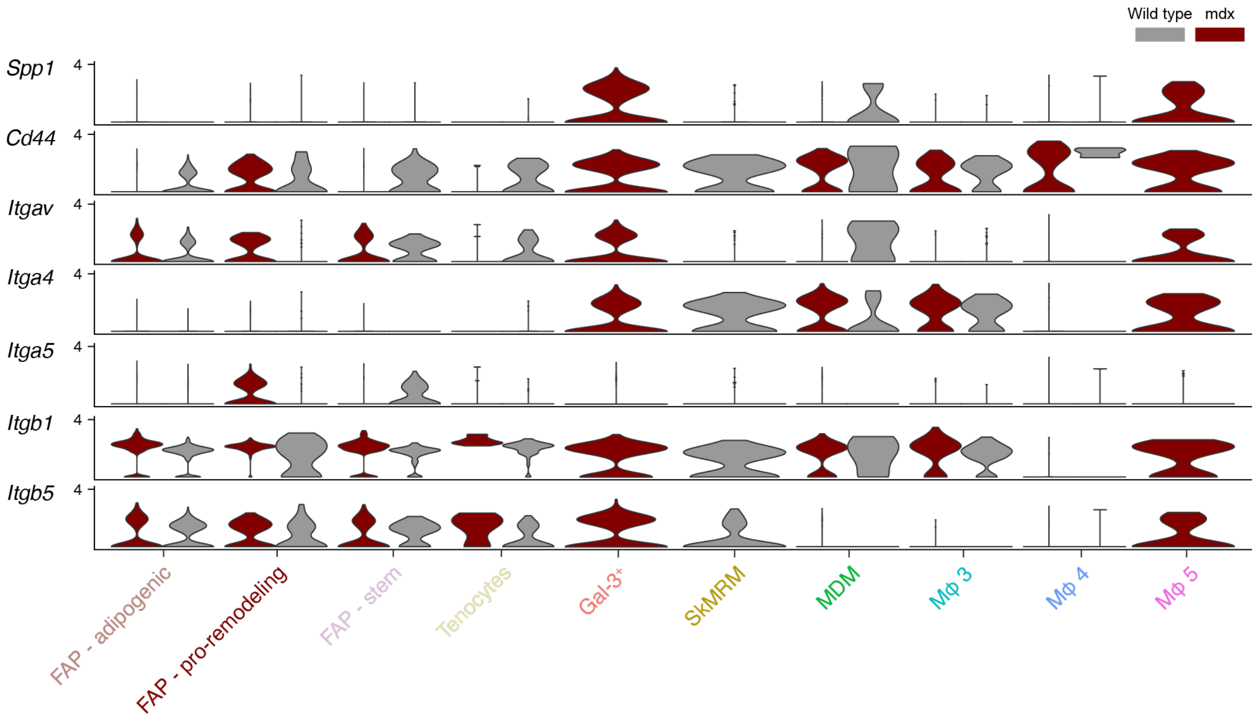


**Fig. S8. Expression of Spp1 and its receptors in WT and mdx FAPs and macrophages.** Violin plots showing the expression of Spp1, CD44 and integrins in macrophages and FAPs from WT (grey) and mdx mice (dark red).

**Supplementary Figure 9**

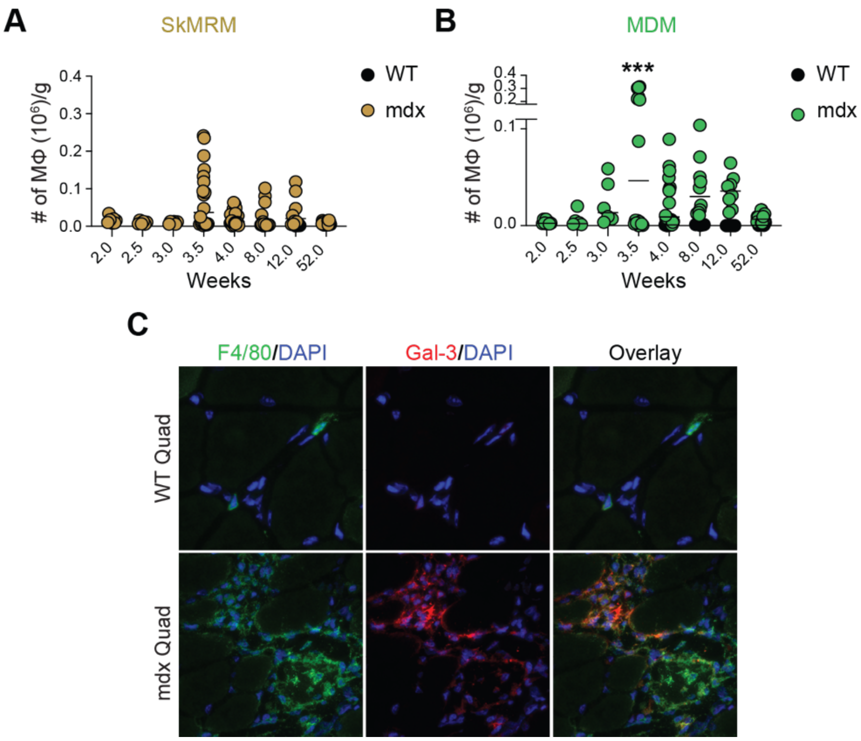


**Fig. S9. Regulation and localization of muscle macrophage populations in muscular dystrophy.** (**A** and **B**) Enumeration of SkMRMs (A) and MDMs (B) during muscular dystrphy. n= 7-9 for each time point. (**C**) Immunofluorescence staining of macrophages in dystrophic muscle. Immunofluorescence localization of Gal-3$^{+}$ Mϕ using a galectin-3-specific antibody (red). F4/80 was used a pan macrophage marker (green). *p<0.05, **p<0.01, ***p<0.001, ****p<0.0001 using a 2-way ANOVA with Sidak's multiple comparison test.

**Supplementary Figure 10**

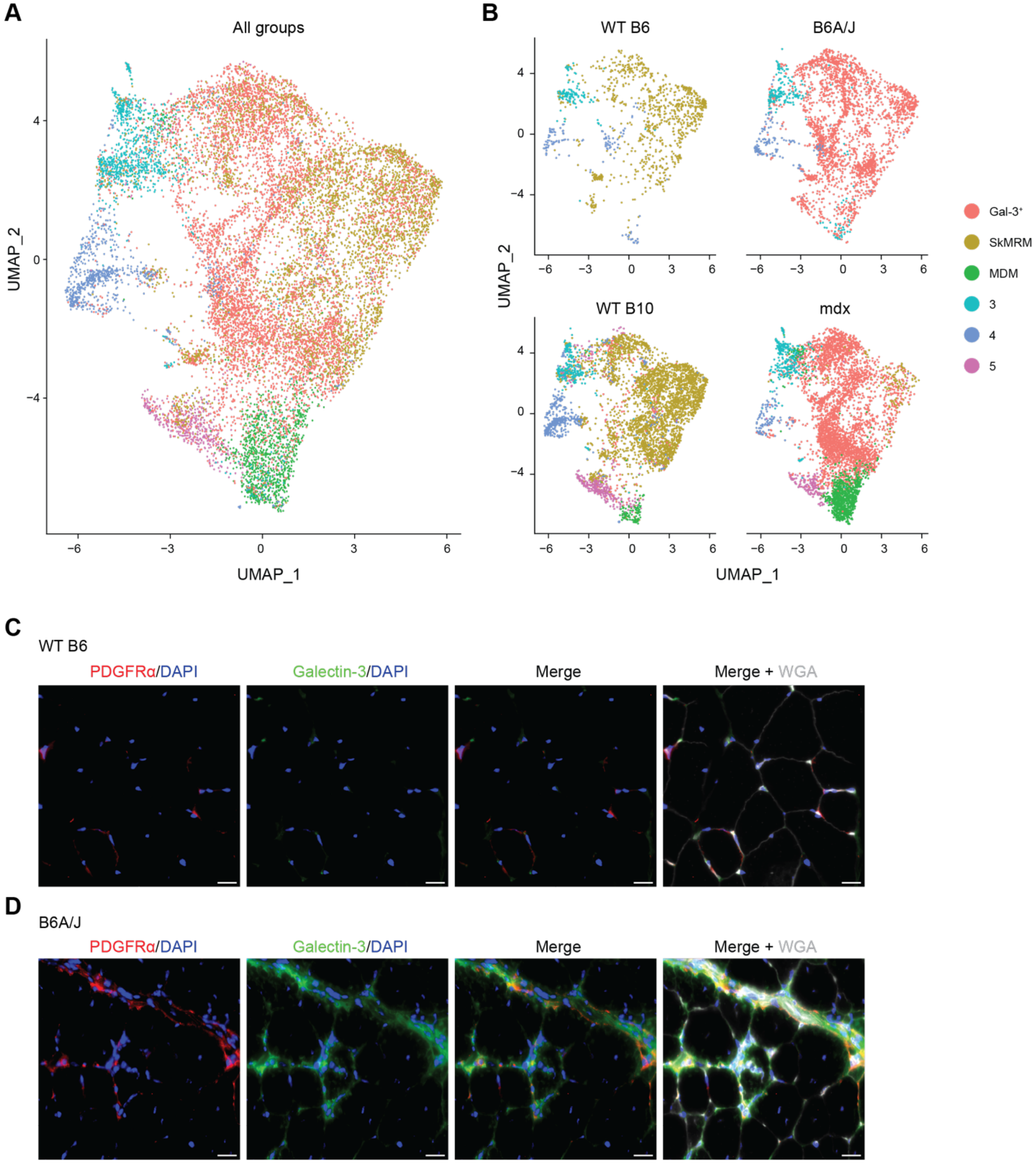


**Fig. S10. Reference-based integration of LGMD2B muscle macrophage scRNAseq datasets.** (**A**) UMAP plot of combined heterogeneous muscle macrophages datasets from mdx (n= 1, 4-wk-old), B6A/J (n= 2, 8-mon-old, pooled) and WT B10 (n= 1, 4-wk-old) and WT B6 (n= 1, 8-mon-old) control mice. (**B**) UMAP plots for each genotype showed a predominant muscle macrophage subset corresponding to Gal-$3^+$ macrophages in B6A/J while cluster 1 (SkMRM) is the dominant population in the matching B6 control mice. (**C** and **D**) Immunofluorescence staining of 12-month-old B6 (C) B6A/J muscle (D) with PDGFR$\alpha$ (red) and galectin-3 antibodies (green). Frozen sections were counterstained with DAPI and wheat germ agglutinin (WGA) to highlight nuclei and the ECM, respectively.

### Supplementary Figure 11

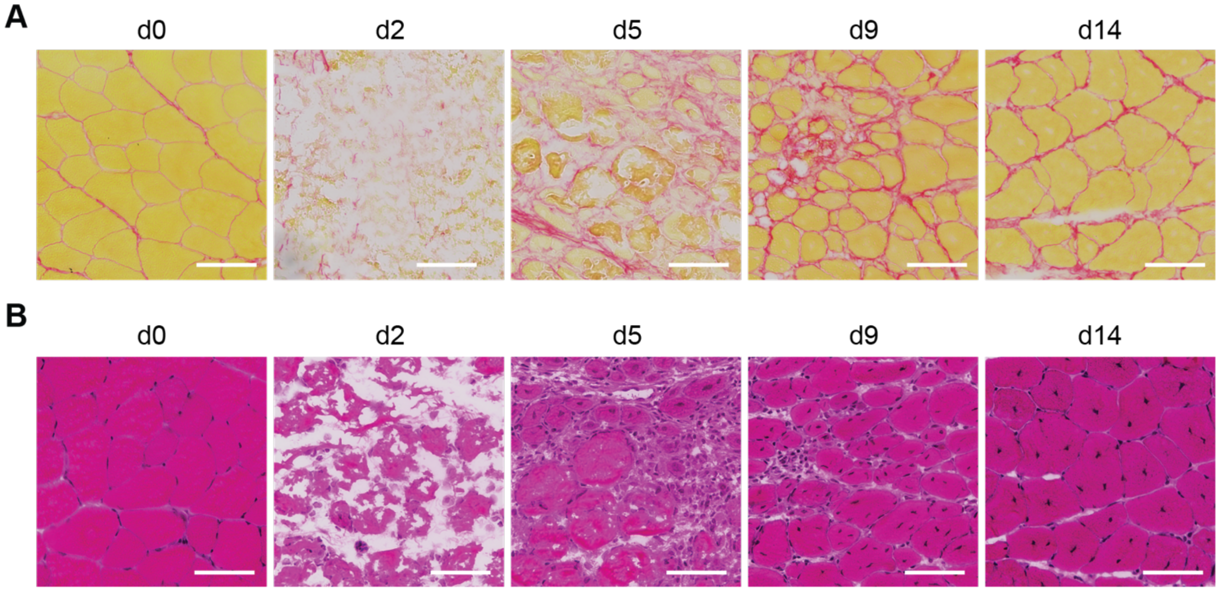


**Fig. S11. Histological examination of acutely-injured muscle.** (**A, B**) Muscle histology at different time points following $BaCl_2$-induced acute injury in 6-weeks old B6 mice. Representative sirius red (A) and Hematoxylin and eosin (H&E) staining (B) of quadriceps cross sections.

**Supplementary Figure 12**

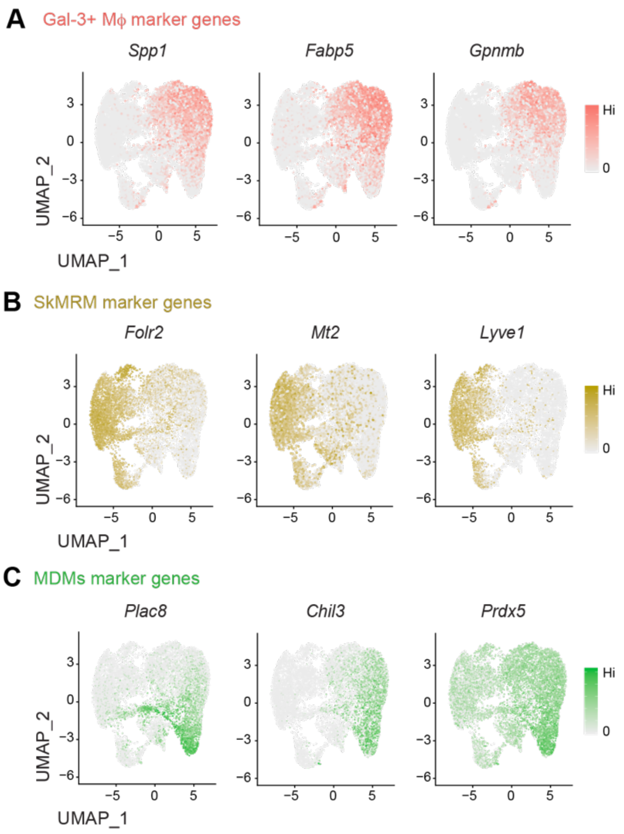


**Fig. S12. Preferential expression of Gal-3⁺ Mϕ, SkMRMs and MDMs marker genes.** (A-C) Feature plots of macrophage marker genes in Gal-3⁺ Mϕ, (Spp1, Fabp5, Gpnmb), SkMRM (Folr2, Mt2, Lyve1) and MDMs (Plac8, Chil3, Prdx5).

1. S. N. Oprescu, F. Yue, J. Qiu, L. F. Brito, S. Kuang, Temporal Dynamics and Heterogeneity of Cell Populations during Skeletal Muscle Regeneration. *iScience* **23**, 100993 (2020).
2. T. Stuart, A. Butler, P. Hoffman, C. Hafemeister, E. Papalexi, W. M. Mauck, 3rd, Y. Hao, M. Stoeckius, P. Smibert, R. Satija, Comprehensive Integration of Single-Cell Data. *Cell* **177**, 1888-1902 e1821 (2019).
3. C. Hafemeister, R. Satija, Normalization and variance stabilization of single-cell RNA-seq data using regularized negative binomial regression. *Genome Biol* **20**, 296 (2019).
4. Y. Hao, S. Hao, E. Andersen-Nissen, W. M. Mauck, 3rd, S. Zheng, A. Butler, M. J. Lee, A. J. Wilk, C. Darby, M. Zager, P. Hoffman, M. Stoeckius, E. Papalexi, E. P. Mimitou, J. Jain, A. Srivastava, T. Stuart, L. M. Fleming, B. Yeung, A. J. Rogers, J. M. McElrath, C. A. Blish, R. Gottardo, P. Smibert, R. Satija, Integrated analysis of multimodal single-cell data. *Cell* **184**, 3573-3587 e3529 (2021).
5. S. Jin, C. F. Guerrero-Juarez, L. Zhang, I. Chang, R. Ramos, C. H. Kuan, P. Myung, M. V. Plikus, Q. Nie, Inference and analysis of cell-cell communication using CellChat. *Nat Commun* **12**, 1088 (2021).
6. A. Crotti, H. R. Sait, K. M. McAvoy, K. Estrada, A. Ergun, S. Szak, G. Marsh, L. Jandreski, M. Peterson, T. L. Reynolds, I. Dalkilic-Liddle, A. Cameron, E. Cahir-McFarland, R. M. Ransohoff, BIN1 favors the spreading of Tau via extracellular vesicles. *Sci Rep* **9**, 9477 (2019).
7. C. P. Hans, N. Sharma, S. Sen, S. Zeng, R. Dev, Y. Jiang, A. Mahajan, T. Joshi, Transcriptomics Analysis Reveals New Insights into the Roles of Notch1 Signaling on Macrophage Polarization. *Sci Rep* **9**, 7999 (2019).
8. V. A. Schneider, T. Graves-Lindsay, K. Howe, N. Bouk, H. C. Chen, P. A. Kitts, T. D. Murphy, K. D. Pruitt, F. Thibaud-Nissen, D. Albracht, R. S. Fulton, M. Kremitzki, V. Magrini, C. Markovic, S. McGrath, K. M. Steinberg, K. Auger, W. Chow, J. Collins, G. Harden, T. Hubbard, S. Pelan, J. T. Simpson, G. Threadgold, J. Torrance, J. M. Wood, L. Clarke, S. Koren, M. Boitano, P. Peluso, H. Li, C. S. Chin, A. M. Phillippy, R. Durbin, R. K. Wilson, P. Flicek, E. E. Eichler, D. M. Church, Evaluation of GRCh38 and de novo haploid genome assemblies demonstrates the enduring quality of the reference assembly. *Genome Res* **27**, 849-864 (2017).
9. C. Trapnell, L. Pachter, S. L. Salzberg, TopHat: discovering splice junctions with RNA-Seq. *Bioinformatics* **25**, 1105-1111 (2009).
10. A. Frankish, M. Diekhans, A. M. Ferreira, R. Johnson, I. Jungreis, J. Loveland, J. M. Mudge, C. Sisu, J. Wright, J. Armstrong, I. Barnes, A. Berry, A. Bignell, S. Carbonell Sala, J. Chrast, F. Cunningham, T. Di Domenico, S. Donaldson, I. T. Fiddes, C. Garcia Giron, J. M. Gonzalez, T. Grego, M. Hardy, T. Hourlier, T. Hunt, O. G. Izuogu, J. Lagarde, F. J. Martin, L. Martinez, S. Mohanan, P. Muir, F. C. P. Navarro, A. Parker, B. Pei, F. Pozo, M. Ruffier, B. M. Schmitt, E. Stapleton, M. M. Suner, I. Sycheva, B. Uszczynska-Ratajczak, J. Xu, A. Yates, D. Zerbino, Y. Zhang, B. Aken, J. S. Choudhary, M. Gerstein, R. Guigo, T. J. P. Hubbard, M. Kellis, B. Paten, A. Reymond, M. L. Tress, P. Flicek, GENCODE reference annotation for the human and mouse genomes. *Nucleic Acids Res* **47**, D766-D773 (2019).
